# Dynamics in the Intact fd Bacteriophage Revealed by Pseudo 3D REDOR-Based Magic Angle Spinning NMR

**DOI:** 10.1101/2024.06.17.598843

**Authors:** Orr Simon Lusky, Dvir Sherer, Amir Goldbourt

**Affiliations:** School of Chemistry, Faculty of Exact sciences, Tel Aviv University, Tel Aviv 6997801, Israel; The Center for Physics and Chemistry of Living Systems, Tel Aviv University, Tel Aviv 6997801, Israel

**Keywords:** magic-angle spinning, solid-state NMR, protein dynamics, filamentous bacteriophage, order parameter, dipolar coupling

## Abstract

The development of robust NMR methodologies to probe dynamics on the atomic scale is vital to elucidate the close relations between structure, motion, and function in biological systems. Here we present an automated protocol to measure, using magic-angle spinning NMR, the effective ^13^C-^15^N dipolar coupling constants between multiple spin pairs simultaneously with high accuracy. We use the experimental dipolar coupling constants to quantify the order parameters of multiple C-N bonds in the thousands of identical copies of the coat protein in intact fd-Y21M filamentous bacteriophage virus, and describe its overall dynamics on the sub-millisecond time scale. The method is based on combining three pseudo three-dimensional NMR experiments, where a rotational echo double resonance (REDOR) dephasing block, designed to measure internuclear distances, is combined with three complementary ^13^C-^13^C mixing schemes: dipolar-assisted rotational resonance, through-bond transfer-based double quantum / single quantum correlation, and radio-frequency driven recoupling. These mixing schemes result in highly resolved carbon spectra with correlations that are created by different transfer mechanisms.

We show that the helical part of the coat protein undergoes a uniform small (∼30°) amplitude motion, while the N-terminus is highly flexible. In addition, our results suggest that the reduced mobility of lysine sidechains at the C-terminus are a signature of binding to the single stranded DNA.

## Introduction

Dynamic fluctuations of biomolecules are fundamental to their function. Examples include enzymatic catalysis and self-assembly processes that necessitate mobility of essential constituents. Elucidating such dynamic behaviours is therefore of great benefit for understanding many biological processes, an example of which is the life cycle of viruses. A virus has to attach to the recognition site in the host cell, inject its genetic material (either DNA or RNA) and the genome must be replicated and translated to generate new viral proteins. Once the proteins are produced, they undergo self-assembly with the genetic material to create new virions capable of infecting new cells. All of these stages necessitate different degrees of molecular flexibility, with variations in the time scales and the amplitudes of motion. Characterising these motions offers invaluable information on the processes that govern viral replication.

Bacteriophages (phages) are viruses that infect bacteria. Filamentous bacteriophages of family *inoviridae* contain a circular ssDNA genome encapsulated by thousands of identical copies of gVIIIp, the major coat protein. The Ff filamentous phages (fd, f1, M13) infect *E. Coli* bearing F-pili.^1^ Their capsid, mainly containing gVIIIp, is arranged from pentamers related by a two-fold screw axis (class-I symmetry) that wrap the ssDNA. Ff phages were thoroughly studied as a model system for understanding phage structure,^2–4^ for studying viral morphogenesis^5,6^ and for soft matter physics.^7,8^ Filamentous phages have also proven useful for various applications such as phage display^9,10^ and electrochemistry^11,12^. Considering the ubiquitous possible usages of these viruses, it is beneficial to establish robust methods to study the property of dynamics in such systems.

Magic Angle Spinning solid-state NMR (ssNMR) has proven to be a powerful tool for studying dynamics in biomolecules. The ability to probe motions at different time scales was extensively reviewed^13–15^ and was demonstrated on various systems such as amyloids,^16,17^ membrane proteins,^18,19^ and viral proteins assemblies.^20–22^ One of the methods to quantify dynamics in molecules is by measuring motionally-averaged anisotropic interactions. The order parameter, *𝒮*, describes the ratio between the breadth of the experimental interaction tensor to its equivalent in the rigid case, when the motion is sufficiently faster than the time scale of the interaction under investigation. Naturally, the values of *𝒮* vary between 1 (rigid limit) and 0 (isotropic motion). When assuming a cylindrical-symmetric motion (“diffusion on a cone”),^23^ the order parameter can be related to the amplitude of motion, θ, according to

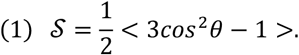

In the “diffusion in a cone” model,^24^ the order parameter is related to the maximal angle θ by

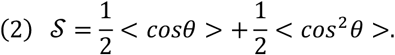

Other types of motions can also be modelled and simulated.^25–27^

The calculation of *𝒮* is straightforward when measuring the motionally averaged dipolar coupling tensor, as the rigid limit is well defined by the direct dependence of the dipolar coupling constant *d* on the inverse cube of the distance 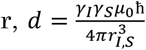 (*γ* and *γ* are the gyromagnetic ratios of spins I and S respectively, and other symbols have their usual meanings). Examples for studying dynamics using averaged dipolar coupling tensors span different types of proteins and their complexes.^28–32^ Early studies utilised mainly the dipolar chemical shift correlation (DIPSHIFT) technique,^33,34^ and other studies utilised various symmetry-based sequences to recouple dipolar couplings.^35–38^ Recently it was shown that DIPSHIFT, and the robust and accurate rotational-echo double resonance (REDOR) technique to measure effective heteronuclear dipolar coupling constants,^39^ are in fact theoretically similar.^40^ In order to implement REDOR two experiments are required – a reference echo experiment (with a signal denoted as ‘S_0_’) and another experiment in which dephasing pulses are employed on a coupled spin (with a signal denoted as ‘S’). The dephasing depends on the dipolar coupling constant and for a system of two half-spins the resulting REDOR recoupling curve (for ideal pulses) follows the analytical expression^41^

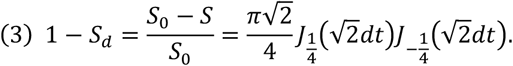

Here ‘*d*’ is the effective dipolar coupling constant and ‘*t*’ is the dephasing time. By performing multiple experiments at different dephasing times, the signal can be fitted to Eq. (3). For small molecules with a well-resolved 1D spectrum, a set of several 1D experiments spanning the initial rise up to 1 − *S*_*d*_ *≈ 0*.*6* is sufficient for obtaining accurate dipolar coupling constants. However, for proteins with hundreds of spins, the addition of more dimensions greatly enhances the spectral resolution. Pseudo three-dimensional (p3D) experiments with various mixing schemes, in which the pseudo-dimension is REDOR, were proven to be useful in calculating distances and probing dynamics in specific sites in biomolecules: ^13^C{^19^F} REDOR with radio-frequency drive recoupling (RFDR) mixing,^42 13^C{^2^H} REDOR with CORD mixing^43^ and 2D heteronuclear-resolved ^13^C{^1^H}, ^15^N{^1^H} REDOR^32^ experiments – all demonstrated on the model protein GB1. The latter was used for measuring the order parameters of most backbone N-H and Cα-H bonds in the fully protonated protein using ^1^H detection.

Generally, in order to observe a reduction in the dipolar interaction, the motion must have a sufficiently large (*≳* 10*°*) amplitude and be at least an order of magnitude faster than the typical time scale of that interaction. As a result, in comparison to measurements of ^1^H-^15^N and ^1^H-^13^C order parameters, the range of motions that can be detected by measuring the effective ^13^C-^15^N dipolar coupling constants is one order of magnitude slower than those involving protons, namely – longer correlation times. Kashefi et. al. applied ^13^C{^15^N} REDOR with DARR mixing to study the average couplings of Cα-N and C-N in the peptide MLF and in the cytoplasmic fragment of the *E. coli* aspartate receptor (CF).^44^ They showed that the REDOR block can be used as a filter for DARR experiment, providing information on the ratio of rigid and dynamic residues. Peak volume analysis of different crosspeaks in this REDOR-DARR experiment reports on this ratio and such an analysis was used to quantify the distribution of mobile residues in CF. Yet, site-specific order parameters have not been derived.

Similar experiments involving a combination of REDOR with various different homonuclear mixing sequences can be used to quantify to high accuracy the individual order parameters of each residue in the protein if the experiments are recorded in a p3D fashion and the build-up curves are site-specifically analysed. The choice of mixing scheme is important for the type of observed correlations. Mixing based on dipolar interaction can show more correlations, as no chemical bond is required, and since the dipolar interaction is generally stronger than the scalar coupling. However, large-amplitude motions average the dipolar interaction, causing the mixing to be less efficient, and with the effects of dipolar truncation, dynamic spins may not appear at sufficient sensitivity after the mixing period. Scalar-based experiments on the other hand are less compromised by dynamic spins due to their longer transverse relaxation times and due to the fact that this interaction is not averaged out by motion. Thus, such experiments can significantly enhance correlations that do not appear in dipolar-based experiments. Experiments with isotropic mixing like RFDR^45^ utilise both the dipolar and the scalar interactions as transfer mechanisms, allowing observation of both rigid and dynamic regions. For example, with RFDR and short mixing times we could clearly observe crosspeaks of residues 1-3 of the fd coat protein.^46^ The INADEQUATE experiment^47^, adapted to ssNMR^48^, is another possible candidate for such studies. The double-quantum (DQ) frequencies observed in the indirect dimension reduce the spectral congestion and facilitate the assignment process. This property, along with its enhanced sensitivity to detect dynamic regions when DQ excitation is based on scalar-coupling, was also shown to aid the assignment of mobile residues in fd coat protein.^49^ Therefore, a combination of REDOR as a dephasing block with INADEQUATE mixing is an adequate candidate for quantitatively studying highly dynamic residues in viruses and other biological macromolecules and complexes. The three mixing schemes are therefore complementary in terms of their information content.

In this study we present the dynamics profile of C-N bonds in the major coat protein of the fd phage possessing a Y21M mutation in gVIIIp. We combined DARR, RFDR, and INADEQUATE mixing schemes with a REDOR dephasing block in order to quantify the amplitudes of motion of Cα-N bonds in the backbone of 36 of its 50 residues (including the highly flexible N-terminus) and of several sidechains containing C-N moieties. We demonstrate that overall, the helix has uniform order parameters, and the N-terminus undergoes large-amplitude motions with an order parameter as low as 0.16. Analysis of sidechain dynamics revealed that the single tryptophan is highly rigid, and that the lysine residues next to the ssDNA exhibit a somewhat restricted motion compared to other sidechains like K8 and Q15, although this motion is of larger amplitudes than the backbone. Finally, we demonstrate the utility of REDOR as a filter to simplify INADEQUATE spectra.

## Materials and Methods

### Sample

A uniformly^-13^C, ^15^N labelled sample of fd-Y21M bacteriophage was prepared using our lab protocols,^50^ and packed into a 4mm ZrO_2_ MAS rotor.

### NMR methods

Magic-angle spinning NMR experiments were conducted on a Bruker Avance III spectrometer operating at 14.1T, equipped with a MAS 4mm E-free probe. The chemical shifts of ^13^C were externally referenced to Adamantane at 40.48ppm.^51^ Experiments were conducted at an estimated temperature of ∼11±1°C, calibrated according to the chemical shift of water protons in the sample. The complete list of all experimental parameters appears in section S1 in the supporting information (SI).

### Data analysis

NMR data were processed using NMRPipe.^52^ The analysis of spectra was done using SPARKY 3.134^53^ and POKY.^54^ RAVEN^55^ was used for crosspeak analysis and for generating build-up curves, which were fit to the universal REDOR curves using a home-written MATLAB script (see section S2 in the SI). Linear prediction was used in the indirect dimension of the REDOR-RFDR and INADEQUATE-REDOR experiments. The validity of obtaining quantitative results when using linear prediction is discussed in section S4 in the SI.

The identification of spin-pairs belonging to different crosspeaks in REDOR-DARR and REDOR-RFDR experiments relied on our prior assignment of the fd-Y21M phage coat protein^50^ (BMRB entry 26910). The assignment table and the list of the crosspeaks from all the spectra were loaded into the RAVEN software and automatically assigned. The assigned crosspeaks were then analysed using the ‘Cross Peaks Analysis Tool’ of RAVEN resulting in a table of intensities, containing S(t) and S_0_(t) for each crosspeak. Several crosspeaks were analysed manually (peaks were identified and the intensity was extracted) in order to validate the automated RAVEN process.

Since the current version of RAVEN does not support peak identification from DQ data, as it relies on the BMRB assignment table, we wrote a Python script as an extension to support scalar-coupling-based double-quantum frequencies thus enabling analysis of our INADEQUATE-REDOR data. The script creates a new table from the original BMRB assignment table that includes all possible ^1^JCC-based DQ resonances. The revised assignment list can then be used as input for both RAVEN and POKY. The script is available for download via Github, https://github.com/amirgoldbourt/DQ-extension-for-RAVEN-and-POKY. A more detailed description of the script appears in section S3 in the SI and demonstrated in the supplementary video POKY and supplementary video RAVEN.

In our analysis of the REDOR recoupling curve, we fit the crosspeak intensities from the homonuclear spectra acquired at different mixing times to the universal curve, Eq. (3), using a home-written MATLAB script (given in section S2 in the SI). This analysis inherently assumes a two-spin system and the application of ideal pulses. In order to check the validity of the first assumption, we used the NMR simulation package SIMPSON^56^ to calculate a REDOR recoupling curve for a three-spin system consisting one ^13^C and two ^15^N spins, with dipolar coupling constants of 1005Hz and 200Hz, corresponding to the typical distances between Cα and the two N spins of the same and the adjacent amino acids. The fit of the simulated data to the universal curve corresponding to an isolated Cα−N spin pair in the same amino acid resulted in a deviation of 2Hz, which is smaller than the experimental error, thus validating the approximation. Similarly, simulating a C_3_N spin system containing homonuclear ^13^C-^13^C dipolar couplings of Cα with C and Cβ resulted in a deviation of 4Hz from the fit to Eq. (3). The results of the simulations are shown in section S4 in the SI. The second assumption ignores the finite pulse correction^57^ (underestimates the dipolar coupling constant by ∼2% for our experimental conditions) when using the universal curve, however, the fit error due to the experimental noise averages ∼5-10% (as derived from χ^2^ fitting) and is higher. This error mainly includes the effect of imperfect π pulses and instability in the spinning speed.

Once all dipolar coupling constants were obtained, the order parameter was calculated as

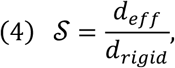

with *d*_*rigid*_ being the dipolar coupling constant at the rigid limit, 1005 Hz, determined by the bond length of Cα-N (1.45Å).^58^ An example of the data analysis appears in the section S5 in the SI.

## Results and Discussion

The general scheme of the experiments, shown in Figure 1, includes a 2D ^13^C-^13^C homonuclear mixing sequence along with a REDOR dephasing block. Three mixing schemes were used: DARR, RFDR and INADEQUATE. The DARR and RFDR mixing times were chosen to be 5 ms and 6 ms respectively to limit the magnetisation transfers to spins close in space (*≲* 3Å), decreasing the total number of crosspeaks and thus reducing ambiguity and increasing crosspeak sensitivity. An alternative approach, when the signal to noise ratio is large, is to use long mixing times, thereby generating multiple redundant crosspeaks bearing the same information. In contrast to the DARR and RFDR-based experiments, where the dephasing was in the indirect dimension, the dephasing in the INADEQUATE-based experiment was in the direct dimension to prevent a more complex spin behaviour resulting from dipolar dephasing of double-quantum states. Such an experiment was shown suitable for the measurement of torsion angles in rigid systems.^59^ When combining crosspeak assignments from all three experiments, we were able to resolve and analyse 36 (of the total of 50) residues representing all regions of the coat protein. The remaining 14 residues were not assigned due to spectral overlap in the mostly-helical coat protein. As expected, crosspeaks of the residues belonging to the non-structured, dynamic N-terminus either did not appear in the DARR-based experiment at the short mixing time of 5 ms, or had a very low SNR compared to the RFDR- and INADEQUATE-based experiments (Figure 2). On the other hand, peaks belonging to the backbone of residues in the helical part (residues 6 onwards) had excellent SNR in the DARR spectra.

**Figure 1.**
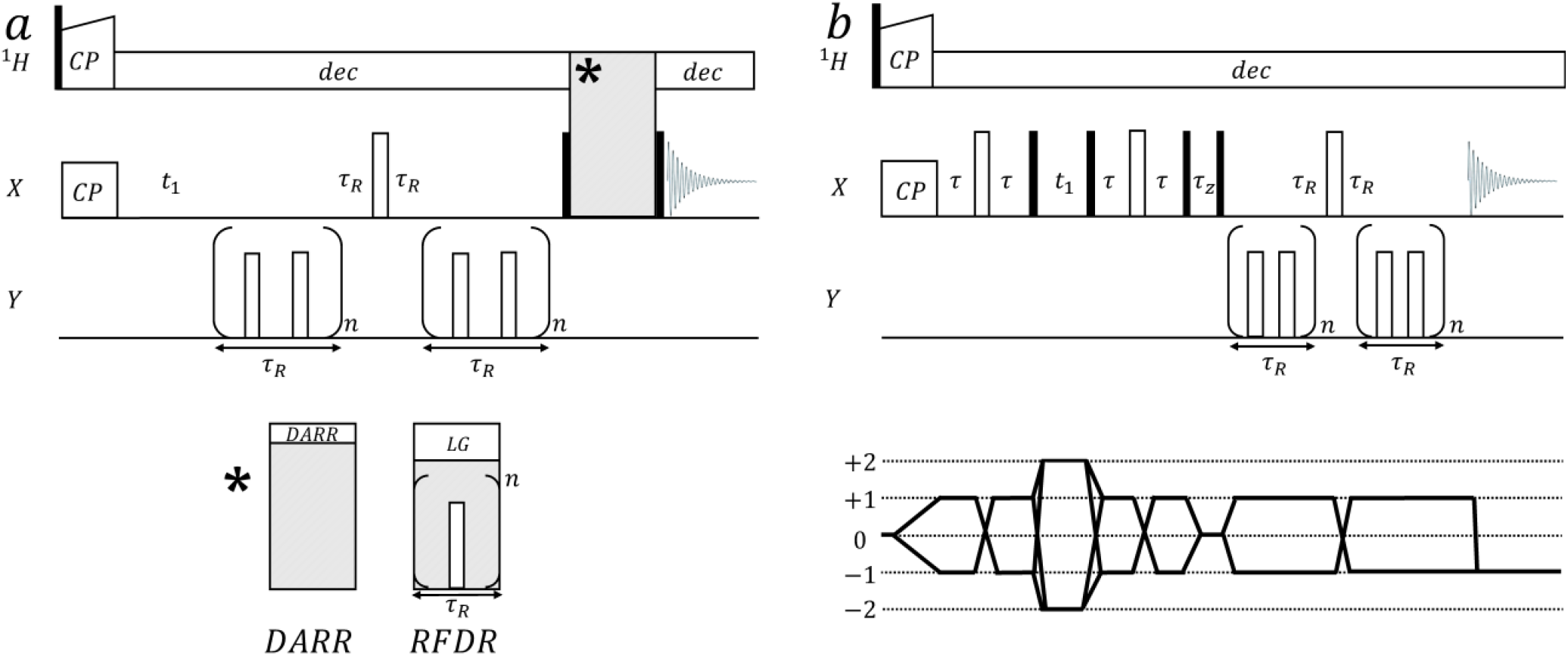
Pseudo 3D pulse sequences for the measurement of 2D-resolved N-C dipolar couplings. Black and white rectangles represent 90° and 180° pulses respectively. CP stands for Cross Polarisation block, ‘dec’ stands for ^1^H decoupling and τ_*R*_ is the MAS rotor period. (a) X{Y}-X REDOR-DARR and X{Y}-X REDOR-RFDR pulse sequences. The two applied schemes are identical with the exception of the mixing block represented by the grey rectangle. For DARR, a ^1^H radio-frequency irradiation with a strength of ν_1_ is applied during mixing at the rotary resonance condition ν_1_ = τ_*R*_^−1^. For RFDR, rotor-synchronous 180° pulses are applied on the X channel (^13^C in our experiments) and a homonuclear ^1^H Lee-Goldburg decoupling is employed. (b) X-X{Y} INADEQUATE-REDOR sequence. The delay time τ was optimised for maximal transfer, τ_*z*_ is the z-filter duration to eliminate non-longitudinal terms. The coherence transfer pathway diagram is drawn beneath. The full phase cycling can be found in Figure S1 in the SI.

**Figure 2.**
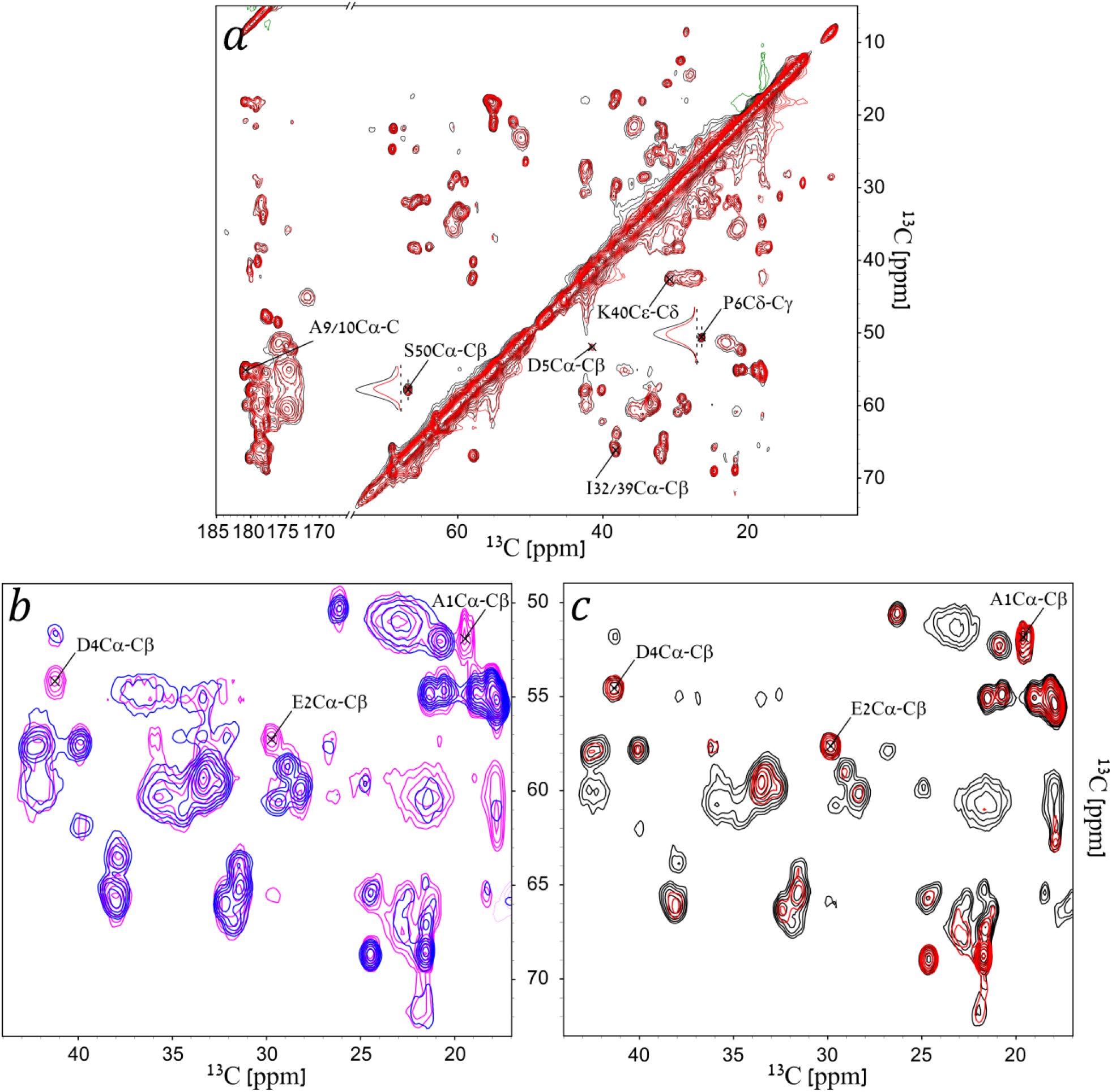
(a) An overlay of 2D REDOR-DARR spectra of gVIIIp in fd-y21m phage – ‘S_0_’ in black and ‘S’ in red. The REDOR dephasing time here is 571μs. Contour levels are drawn at multiples of 1.4 with the lowest matching an SNR of 6. Green represents negative signals. The labels show the assignment for a few crosspeaks that were mentioned in this article. 1D traces of two crosspeaks (P6Cδ-Cγ, and S50Cα-Cβ) are drawn to demonstrate the dephasing on the indirect dimension. (b) An overlay of 2D REDOR-DARR (magenta) and REDOR-RFDR (blue) with a REDOR dephasing time of 429μs demonstrating the emergence of dynamic residues in RFDR. (c) An overlay of 2D REDOR-RFDR ‘S_0_’ (black) and ‘S’ (red) spectra with a REDOR dephasing time of 1143μs. In (b)-(c) spectra contour levels are drawn at multiples of 1.4 with the lowest matching an SNR of 10. The labels show the assignment for the N-terminus residues A1, E2 and D4 that were not detected with DARR mixing.

Figure 2 shows an overlay of two typical 2D DARR-resolved, and two 2D RFDR-resolved spectra comparing ‘S_0_’ and ‘S’. It demonstrates reduction in signal intensities (red spectrum) for crosspeaks originating from Cα (bottom part of the spectrum) while signal intensities of sidechains are retained since they are not sufficiently close to the ^15^N amide to produce dephasing after 571μs (8τ_r_). REDOR curves were then generated from these crosspeaks as detailed in section S5 in the SI. A summary of all build-up curves is given in section S6 in the SI.

### The overall C-N bond dynamics in fd-Y21M coat protein

A total of 142 non-ambiguous and 6 ambiguous (with ambiguity of 2) crosspeaks that involve ^13^C spins dephased by ^15^N, from all p3D spectra, were assigned and fit to the REDOR universal curve. Most fits show good agreement to Eq. (3) with values of 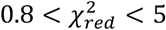. These values show that the assumptions we made, a two-spin system subjected to motions that are faster than the size of the dipolar interaction strength, are acceptable. For the ambiguous crosspeaks (A9-A10, D12-L14, and I32-I39), the fit to a single dephasing curve suggests that the motion of the Cα-N bonds of both amino acids has a similar order parameter. This was also verified by the results of the fit of their corresponding resolved crosspeaks (see section S6 in the SI), and is also supported by our previous observations in which the CSA of the backbone spins shared a similar breadth.^55,60^ The experimental order parameters of the backbone Cα-N bonds are plotted in Figure 3a. This plot shows a trend in which the first residues belonging to the N-terminus (residues 1-5) have larger amplitudes of motion, with an increasing order parameter until reaching a plateau. These results are in agreement with previous reports that showed a flexible non-structured N-terminus.^61,62^ The majority of the protein (residues 6-50) is in a well-defined helix. The average order parameter of the Cα-N bond in the helix is 0.66 ± 0.02 and is lower than order parameters obtained from C-H and even N-H solid-state NMR measurements of other systems such as Pf1 phage coat protein (C-H 0.99 ± 0.04),^63^ GB1 (C-H 0.94 ± 0.04, N-H 0.90 ± 0.07)^64^ and ubiquitin (C-H 0.78 ± 0.09).^30^ When motionally-averaged CSA values were measured,^55,60^ we obtained for the helical part an averaged value of ∼12.5 kHz for C’ and ∼6.3 kHz for the ^15^N amide, very similar to GB1,^65,66^ which is considered to be highly rigid. The order parameters we measure here for Cα-N bonds imply dynamics on a slower time scale due to a smaller anisotropy of the Cα-N (∼1 kHz) dipolar coupling constant compared to the CSA.

**Figure 3.**
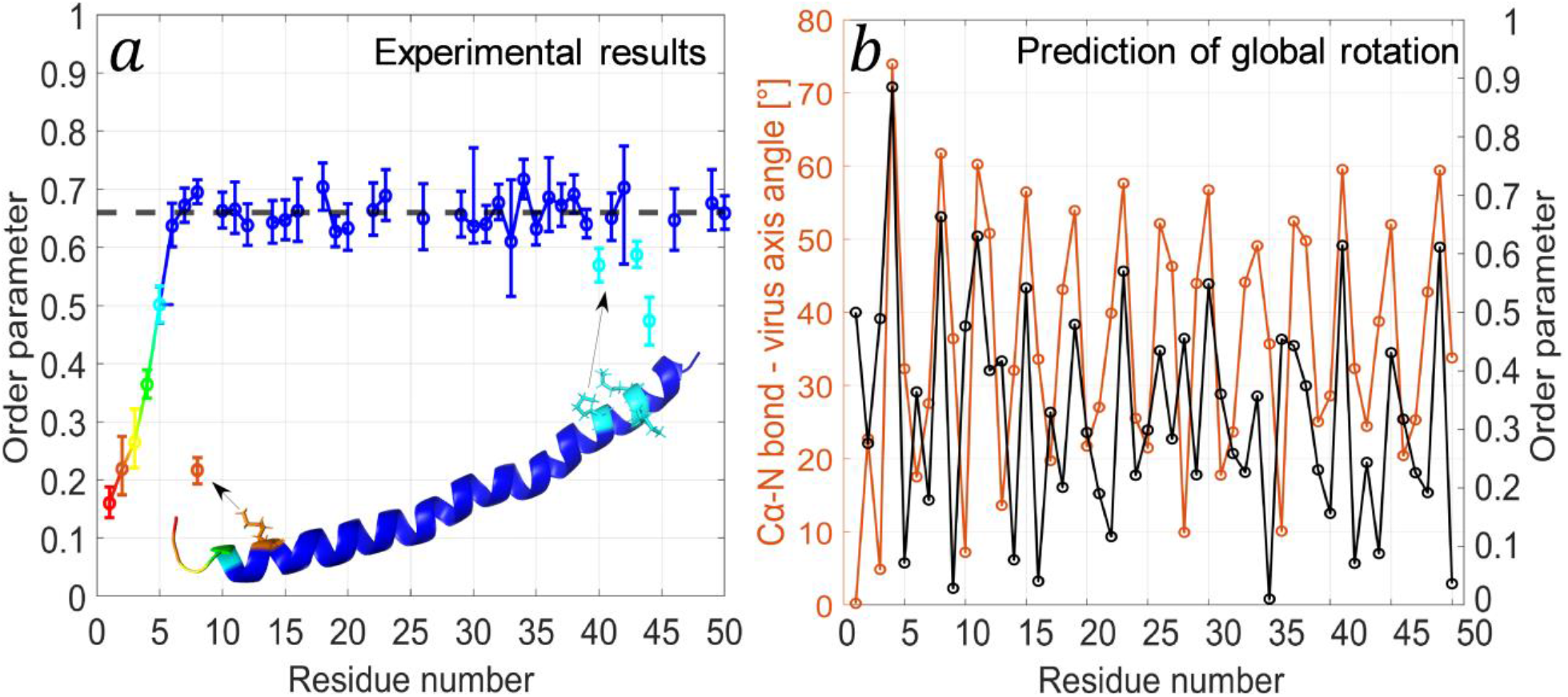
(a) A plot of the order parameters, *𝒮*, of the Cα-N and Cε-N bonds of all residues that appear in at least one of the three types of the p3D REDOR-dephased experiments. The colour gradient (red to blue) represents an increasing order parameter. A cartoon of the known structure^2^ (PDB entry: 2C0X) shows a map of the order parameters with the same colour scheme. In addition, four lysine residues (8, 40, 43, 44) are shown as sticks to represent the order parameter of their Cε-N bond. The black dashed line represents the average order parameter (0.66±0.02) for residues 6-50, and solid lines represent consecutive residues. (b) A plot of the calculated angles (orange) between the viral filament axis and the Cα-N bond vectors for all residues of one coat protein, derived from the known structure^2^ (PDB entry: 2C0X). In black, predicted order parameters matching the Cα-N angles, calculated according to a motion on a cone model (Eq. (1)). Since the experimental method is insensitive to the sign of the interaction, the absolute value of the order parameters was plotted. The results show that rotational motion around the viral axis on a sub-ms time scale is not consistent with our data.

A plausible global motion is a rotation of the entire virus about the axis of the filament, since the virus in its precipitated form is highly hydrated and forms liquid crystals. Such a rotation implies that the Cα-N bonds rotate in a conic motion, which would translate into oscillations in the order parameter. The angles of those bonds relative to the viral axis can be calculated from the known structure,^2^ and in the case of such a global motion, those angles would represent the amplitudes of motion on a cone. However, by calculating the expected order parameters it is clear that the model fails to describe the results. Figure 3b shows the angles between the Cα-N bonds and the vertical axis of the virus, and the absolute value of the calculated order parameters. The calculated periodicity of the order parameters and the large variations in values do not fit the experimental data, and therefore global rotational motion does not occur at a time scale faster than ∼ms.

An alternative option is a uniform motion, namely, a motion with similar amplitudes and overall trajectories that are shared by residues 6-50. In the context of a viral particle, the simplest form of motion that can be suggested is a jittering motion as a result of thermal fluctuations. It is difficult to model such motions and future studies by molecular dynamics, once a plausible model of the ssDNA is obtained, may demonstrate this result.

### Dynamics of the N-terminus

Figure 4a shows the best-fit Cα{^15^N}-CX REDOR build-up curves of residues belonging to the N-terminus. Their values and amplitudes of motion are given in Table 1. Due to the ability of the RFDR experiment to resolve highly dynamic residues, we were even able to observe a quantitative recoupling curve for A1 yielding a very low order parameter of 0.16. As also shown in Figure 3, it is clear that the non-structured N-terminus exhibits large-amplitude motions compared to the rest of the protein. The monotonic increase in the order parameter, hence in the amplitudes of motion, suggests a gradual change, where the tip exhibits the largest motion. The N-terminus in the structure has a shape of a hook that points outwards towards the solvent, and has less inter-residues contacts, possibly allowing the enhanced mobility. REDOR-RFDR spectra showing N-terminus crosspeaks appear in Figure 2.

**Table 1.** Order parameters of the Cα-N bonds in the N-terminus of the fd-Y21M coat protein and the overall average of residues 6-50 as a comparison. The rigid limit was set to be *d*_*rigid*_ = 1005 *Hz*, matching a bond length of 1.45Å. The amplitudes of motion were calculated according to Eqs. (1) and (2).

| Residue | Order parameter | Amplitude (motion on a cone) ($^{\circ}$) | Largest Amplitude (motion in a cone) ($^{\circ}$) |
| --- | --- | --- | --- |
| A1 | $0.16^{+0.02}_{-0.03}$ | $48.4^{+1.2}_{-1.4}$ | $75.2^{+2.0}_{-2.2}$ |
| E2 | $0.22^{+0.04}_{-0.06}$ | $46.2^{+1.7}_{-2.1}$ | $70.8^{+3.3}_{-4.1}$ |
| G3 | $0.26^{+0.04}_{-0.06}$ | $44.4^{+1.7}_{-2.2}$ | $67.5^{+3.1}_{-4.0}$ |
| D4 | $0.36^{+0.02}_{-0.02}$ | $40.6^{+0.9}_{-1.0}$ | $60.7^{+1.6}_{-1.7}$ |
| D5 | $0.50^{+0.03}_{-0.03}$ | $35.2^{+1.2}_{-1.3}$ | $51.7^{+2.0}_{-2.1}$ |
| Average | $0.66 \pm 0.02$ | $28.7^{+2.1}_{-1.8}$ | $41.5^{+3.2}_{-2.8}$ |

**Figure 4.**
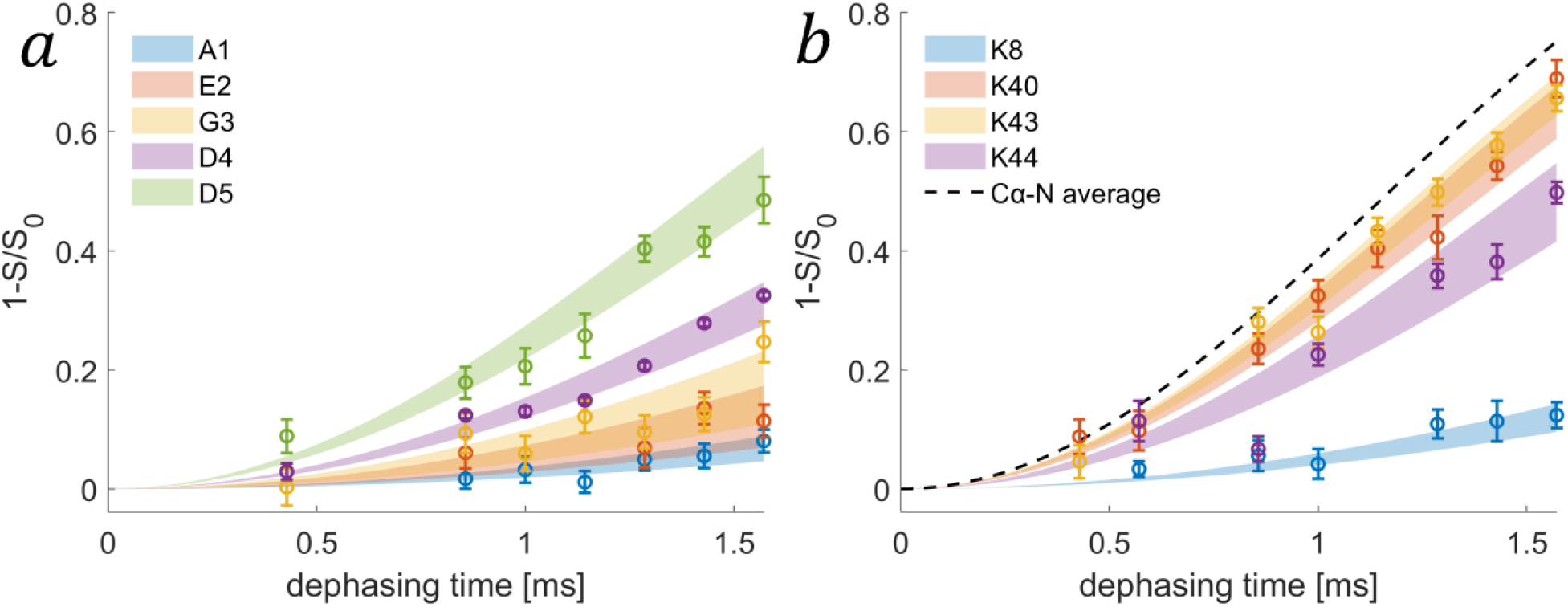
(a) Best-fit ^13^Cα{^15^N}-REDOR build-up curves (from all experimental observations) in the N-terminus residues of fd-Y21M coat protein (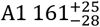 in blue, 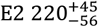 in orange, 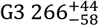 in yellow, 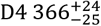in purple, and 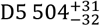 in green). (b) REDOR build-up curves for the average effective dipolar coupling constant calculated for lysine residues (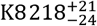in blue, 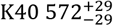 in orange, 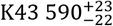 in yellow, and 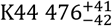 in purple). The dashed curve is the average Cα-N dipolar coupling constant of residues 6-50, 663 Hz. In both (a) and (b) the transparent blocks are the error bars of the fit. The data points are the experimental results, averaged over multiple crosspeaks.

### Dynamics of sidechains

Sidechains of six amino acids contain C-N bonds and therefore their order parameters can also be determined by the proposed p3D methods. They include two positively charged amino acids (R, K) which are commonly involved in binding to negatively charged surfaces such as DNA, or form salt bridges. They also include the polar residues H, Q, and N, and the hydrophobic residue W.

The coat protein of fd-Y21M phage contains five different lysine residues, one tryptophan, and one glutamine. Of the lysines, K8 is located in the start of the helix and is in proximity to the negatively-charged D4 and D5, as well as E20 of a neighbouring coat protein subunit. Other lysines are located in the C-terminus and are in proximity to the ssDNA.^4,67^ The single tryptophan is located in the hydrophobic packing epitope and interacts with other aromatic residues thus stabilising the capsid structure.^3^ Q15 is in the core of the helix.

Table 2 summarises the order parameters of six sidechains (K48 is unresolved). Figure 3a and the REDOR build-up curves shown in Figure 4b, generated by the averaged Cε-Nζ effective dipolar coupling constants of the lysine side chains, show smaller than average order parameters (0.22 for K8, 0.47-0.59 for DNA binding lysines), in agreement with similar observations in Pf1 phage.^30^ One cause for this enhanced dynamic is that the distance between Nζ and the ssDNA allows for water molecules to bridge between the capsid and the nucleic acid.^68,69^ In fact, we have previously observed that lysine to DNA ribose correlations appear only at long mixing times (500ms).^70^ Interestingly we observe that K8 is significantly more dynamic than the other lysine sidechains in contact with DNA although it has three negatively charged residues in its vicinity, D4, D5 and E20. Nevertheless, the distances are probably sufficiently long (4.5-5.9Å) to allow for larger amplitude motions and for the presence of water molecules. These results also suggest that although the lysine-DNA interaction allows for motion despite the electrostatic interactions, it is still more restricted than that of K8, and probably of other sidechains that have motional freedom around the χ_1_, χ_2_, and the other sidechain torsion angles.

**Table 2.** Sidechain order parameters in the fd-Y21M coat protein: Cε-Nζ bond in the lysine residues, Cδ-N_ε2_ in Q15 and C_ε2_-N_ε1_ in W26. The rigid limits for K, Q, and W were set to be *d*_*rigid*_ = 1005*Hz*, 1317*Hz, and* 1116*Hz* respectively, matching the bond lengths of 1.45Å, 1.32Å and 1.40Å.

| #Residue | Order parameter |
| --- | --- |
| K8 | $0.22^{+0.02}_{-0.02}$ |
| Q15 | $0.36^{+0.03}_{-0.03}$ |
| W26 | $0.75^{+0.10}_{-0.08}$ |
| K40 | $0.57^{+0.03}_{-0.03}$ |
| K43 | $0.59^{+0.02}_{-0.04}$ |
| K44 | $0.47^{+0.04}_{-0.04}$ |

The residue W26 was expected to be immobile,^62^ and was even used as a distance reference when solving the structure of the M13 phage.^3^ The order parameter that was obtained from the REDOR curve for the Cε_2_-Nε_1_ bond located in the pyrrole ring has a value of 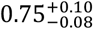, which is higher than any of the other C-N bonds that were detected in all p3D experiments. This fits the previous description of W26 being significantly more rigid due to its role in stabilising the hydrophobic core of the viral capsid.^71^

The order parameter of the sidechain Cδ-Nε_2_ bond in Q15 is relatively low, 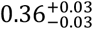, suggesting large amplitude motions. This fits the structure in which that residue points outwards towards the solvent.

### Filtered INADEQUATE

The p3D experiments require the acquisition of multiple REDOR-dephased ‘S’ spectra, including such with dephasing times that cause Cα signal loss of 80% and beyond. Since the dephasing is caused by proximity to ^15^N, the majority of the remaining crosspeaks are those of the sidechains. As a result, these spectra have reduced spectral congestion, and can facilitate the process of assignment and the extraction of many other chemical-shift-based properties. For example, it is highly useful for assigning residues containing carbonyl sidechains (D, E, N and Q) that in many cases overlap with the backbone carbonyl signals. D and E residues are unaffected by the REDOR dephasing pulses, and N and Q sidechain carbonyls tend to be more dynamic compared to the backbone, meaning that the dephasing efficiency of their signal is lower. Indeed, REDOR was used in the past as a filter for DQ-SQ dipolar-based spectroscopy to distinguish aspartate and glutamate from aspartic acid and glutamic acid in ubiquitin and in a membrane protein in lipid bilayers^72^.

We show here that scalar-based filtered INADEQUATE is highly useful, since information about Cα is still maintained in the DQ dimension in the form of crosspeaks with Cβ, meaning that it can still be used for assignment. By choosing the dephasing time according to a theoretical dephasing curve corresponding to the Cα-N distance, the rigid atoms can be filtered out of the spectrum facilitating assignment of resonances that may be unresolved in the non-filtered version.

Figure 5 shows an overlay of the INADEQUATE-REDOR ‘S’ spectrum with the longest dephasing time we recorded, 1714μs, with that of the ‘S_0_’ reference spectrum recorded with a mixing time of 429μs. The upper panel, showing the aliphatic region, demonstrates the selective dephasing of crosspeaks; The sidechains are retained in the ‘S’ spectrum, whereas crosspeaks of Cα undergo dephasing (a site-specific demonstration of threonine residues appears in section S7 in the SI). It is also visible that the signal of the crosspeak corresponding to E2CαCβ-Cα only partially dephased due to elevated dynamics compared to the other similar crosspeaks (e.g., the adjacent E20) demonstrating the potential of an INADEQUATE-REDOR filter as a way to differentiate rigid and dynamic regions. The bottom panel shows the carbonyl region, where the REDOR pulses yielded a more resolved spectrum leaving peaks belonging to sidechain amide and carboxylic groups (E2Cδ, D4Cγ, D5Cγ, Q15Cδ), backbone carbonyl of dynamic residues (G3, D4, D5), and some reduced signals of partially-dephased crosspeaks with similar shifts (G23, G34, G38). One drawback of the experiment is that the long REDOR echo times inherently cause loss of magnetisation due to relaxation. Here an average of 13% decrease in SNR was measured between the reference ‘S_0_’ spectrum (429μs) and ‘S’ spectrum (1714μs) (section S8 in the SI). Yet, this phenomena also occurs in other similar REDOR-filtered experiments,^44,73–76^ relaxation-based filters (T_1_, T_2_, T_1ρ_),^77^ etc.

**Figure 5.**
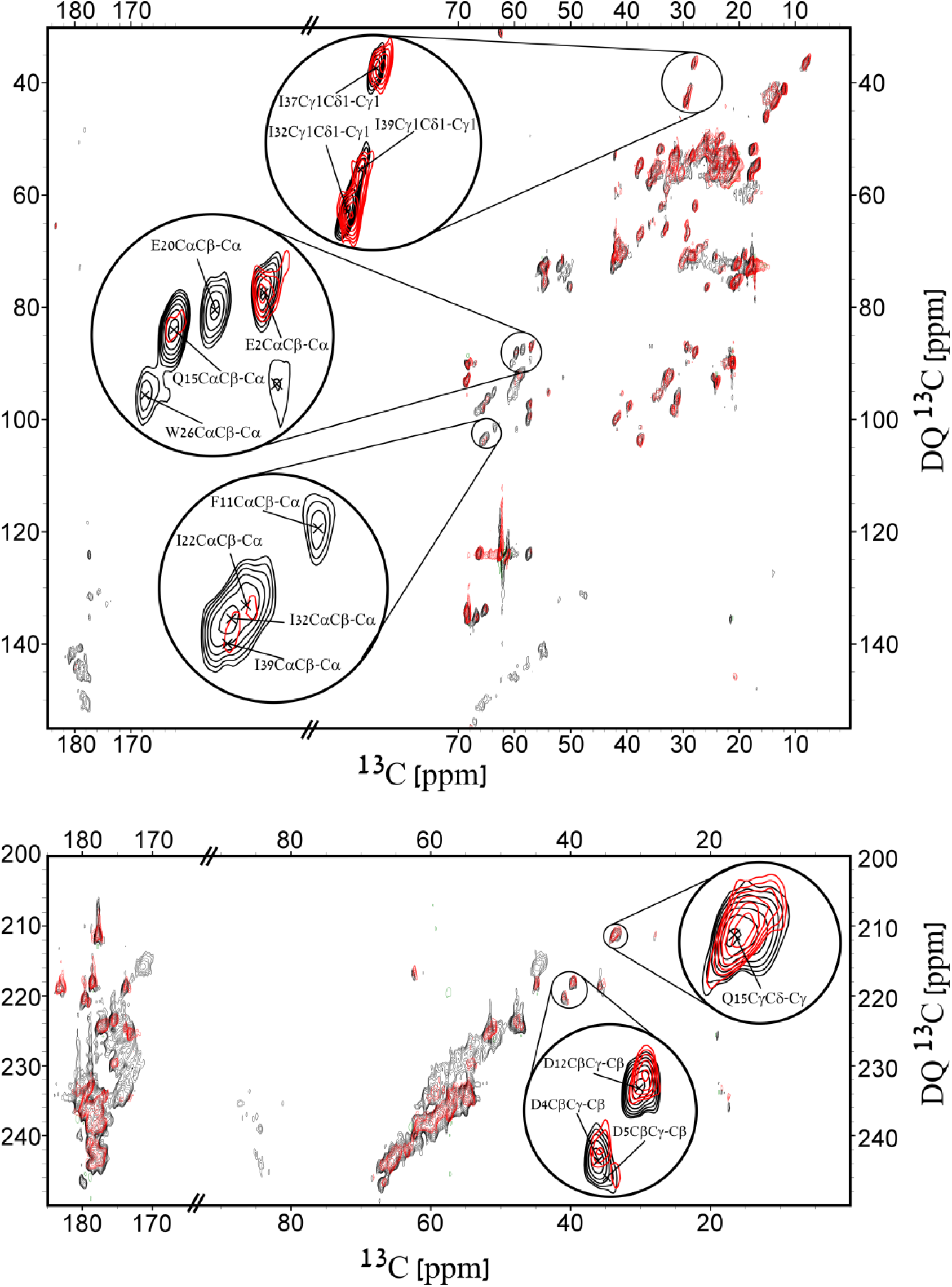
Filtered INADEQUATE-REDOR experiments. The black spectrum is ‘S^0’^ with an echo time of 429μs, and the red is ‘S’ with a dephasing time of 1714μs. The upper and bottom panels show the aliphatic and carbonyl regions of the spectrum respectively, and include magnified crosspeaks in circles. Contour levels are drawn at multiples of 1.3, with the lowest matching an SNR of 6.

## Supporting information

supplementary video POKY

supplementary video RAVEN

## Summary and Conclusions

In this work we presented a set of three pseudo-3D solid-state NMR experiments to study sub-milliseconds dynamics in the coat protein of intact fd-Y21M filamentous bacteriophage virus. The dynamics are characterised by the simultaneous measurement and automated analysis of multiple C-N dipolar order parameters, and the experimental strategies are suitable for many similarly large protein complexes. For the fd coat protein, we show enhanced dynamics of the N-terminus, down to an order parameter of 0.16 for the terminal alanine, in excellent agreement with previous measurements of dynamics that were based on recoupling of the chemical shift anisotropy. The majority of the coat protein was assigned (36/50), with the others being unassigned due to spectral overlap as a consequence of the helical character of the protein. We showed reduced dynamics of ssDNA binding lysine residues with respect to other mobile sidechains, probably resulting from water-mediated electrostatic interactions with the negatively-charged phosphates. The motion of the indole ring of the tryptophan is shown to be restricted, likely due to strong hydrophobic packing interactions. We showed that a global rotational motion of the entire virus along its axis, at the time scale faster than ms, is not in agreement with our data. The uniform distribution of order parameters along the helix implies that the amplitudes of motion at the sub-ms timescale are shared by all the residues in the helix and we hypothesise that such a motion is a result of some uniform jittering. A more accurate description of the motions that lead to the observed order parameters can potentially be predicted from molecular dynamics, and such studies will be possible once a plausible model for the ssDNA is obtained.

The use of three different mixing schemes – DARR, RFDR, and INADEQUATE, was advantageous in the means of getting access to signals originating from both dynamic and rigid spins since they provide complementary spectra that produce correlations based on both dipolar and scalar couplings. The same protocol can be easily generalised by choosing other mixing schemes tailored to the necessity of the system. In some cases, cross-polarisation (CP) may not be efficient for the excitation of dynamic regions even when using long mixing times. In those cases, scalar-based transfer and even direct excitation may be more suitable, however, if the ^1^H-^13^C dipolar coupling is too small to generate polarisation via CP, it is highly likely that the effective ^13^C-^15^N dipolar coupling constant is too small to detect with REDOR and the motion is probably close to isotropic. For the fd-Y21M bacteriophage in this study CP was a competent polarisation transfer mechanism and all dynamic carbons have been efficiently detected.

Finally, we demonstrated the usefulness of REDOR as a filter for INADEQUATE spectra yielding better resolved spectra in the backbone region. The advantage of the DQ approach here is that information on Cα peaks still exists in CαCβ-Cβ crosspeaks that do not undergo dephasing, unlike SQ-based correlations, where the result is the remaining of one side of the diagonal with a similar density of signals.

While demonstrating our experimental approach on an intact filamentous bacteriophage virus, the method can be easily implemented in the study of other viruses and of additional biomacromolecules such as membrane proteins and amyloids. The ability to probe the dynamics of the positively charged amino acids, all of which include nitrogen atoms in their sidechains, is particularly attractive for studying the properties of bio-complexes where negatively-charged protein complexes are common (DNA/RNA, membranes et cetera). In addition, the hurdles of spectral congestion can be overcome for example by using sparsely labelled samples or by resolving the residues of interest by adding additional dimensions. We believe that due to the robustness of this protocol it will be broadly applicable for diverse systems.

## Supporting Information

Includes NMR experimental parameters, matlab scripts for fitting the REDOR curves, description of the double-quantum (DQ) extension for RAVEN and POKY software packages, testing of linear prediction validity in quantitative analysis of peak intensities, all dephasing curves and fits to effective dipolar couplings including error analysis, filtered inadequate experimental examples, two movies demonstrating the use of the DQ extension for RAVEN and POKY.

## Acknowledgements

This research was supported by the Israel Science Foundation, grants #847/17 and #1011/23. We would like to thank Dr. Gili Abramov for the preparation of the fd-y21m phage sample, and Dr. Albert A. Smith-Penzel for illuminating insights.

---

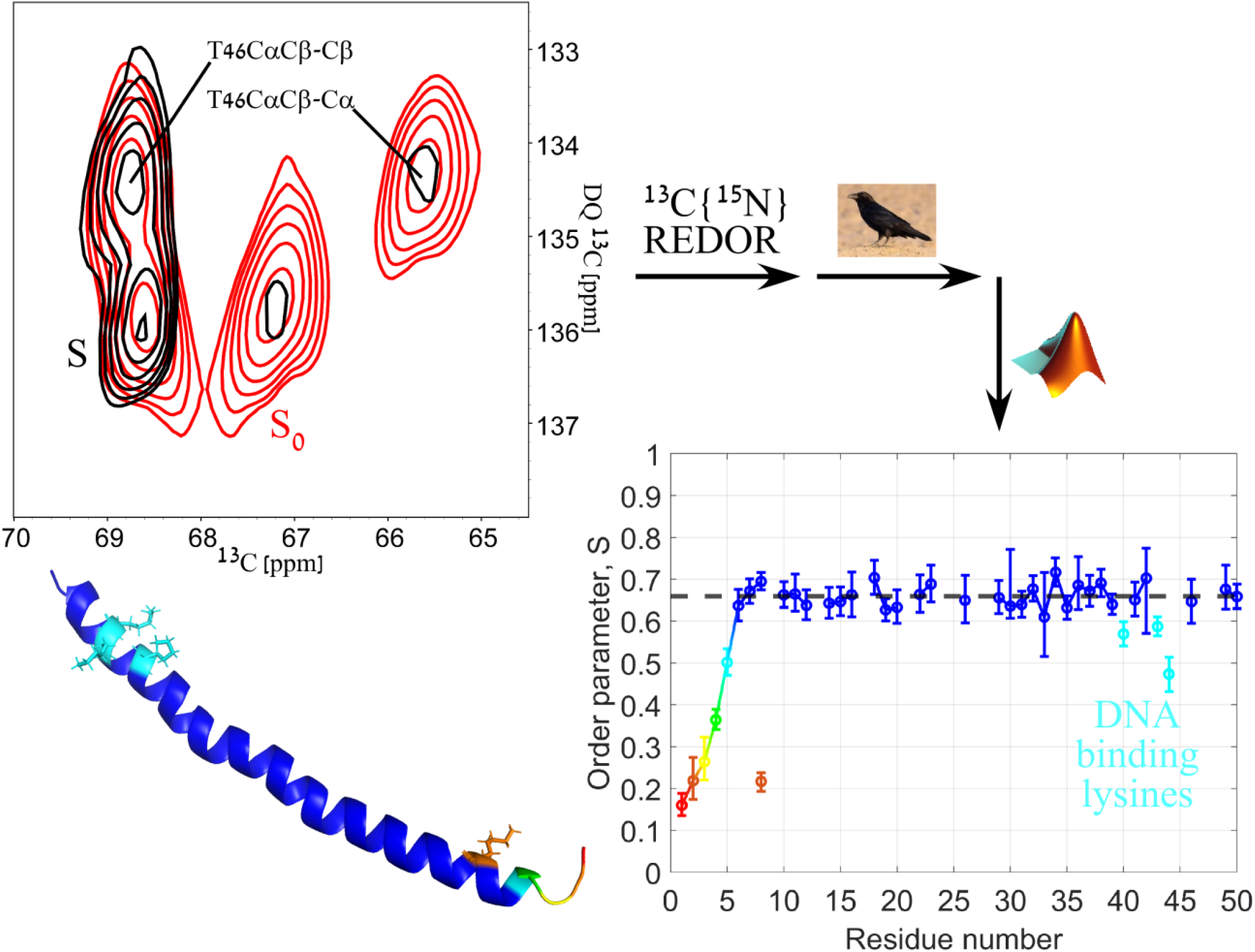

