## Supporting Information for "Dynamics in the Intact fd Bacteriophage Revealed by Pseudo 3D REDOR-Based Magic Angle Spinning NMR"

### S1 Experimental parameters

We conducted three types of experiments, all involving a REDOR dephasing block, and three different mixing schemes: DARR, RFDR and INADEQUATE. In DARR and RFDR the mixing scheme follows the REDOR dephasing, whereas the INADEQUATE mixing precedes the dephasing. All experiments share these parameters:

| | |
|---|---|
| Field [T] | 14.1T |
| Probe | 4mm E-free |
| MAS rate ($\nu_R$) [kHz] | 14.0 |
| Carrier frequency [ppm] | 98.0 |
| $^{1}$H 90° pulse duration [μs] | 2.60 |
| $^{13}$C 90° pulse duration [μs] | 4.50 |
| $^{15}$N 180° pulse duration [μs] | 10.25 |
| CP power ($\nu_H$) [kHz] | 78.5 |
| CP power ($\nu_C$) [kHz] | 55.5 |
| CP contact time [ms] | 1.50 |
| $^{1}$H decoupling power [kHz] | 85 |
| swf-tppm decoupling pulse duration [μs] | 6.0 |
| Relaxation delay (~5$T_1$) [s] | 3.0 |
| Set temperature at the controller [°C] | -15 |
| Estimated actual temperature* [°C] | 11±1 |
| **Processing parameters F1/F2** | |
| Processing software | NMRPipe |
| Zero fill (t1/t2) | 2048/8192 |
| Apodisation (F1/F2) | Squared cosine/Cosine |

*The actual temperature was estimated using the chemical shift of water in the sample.

DARR-REDOR experimental parameters

| | |
|---|---|
| Dephasing times [μs] | 429, 571, 857, 1000, 1143, 1286, 1571 |
| Acquisition mode in indirect dimension | States |
| Acquisition points (t1/t2) | 720/4990 |
| Acquisition times (t1/t2) [ms] | 9.00/25.00 |
| Spectral width F1/F2 [kHz] | 40/100 |

| | |
|---|---|
| $^{1}H$ power during DARR [kHz] | 14 |
| DARR mixing time [ms] | 5.00 |
| Scans | 16 |
| Experimental time ($S_0$ and S) [hours] | ~19 |
| Analysis software | SPARKY 3.134[1] |

RFDR-REDOR experimental parameters

| | |
|---|---|
| Dephasing times [μs] | 429, 857, 1000, 1143, 1286, 1429, 1571 |
| Acquisition mode in indirect dimension | TPPI |
| Acquisition points (t1/t2) | 720/4990 |
| Acquisition times (t1/t2) [ms] | 9.00/25.00 |
| Spectral width F1/F2 [kHz] | 40/100 |
| RFDR mixing time [ms] | 6.00 |
| RFDR $^{13}C$ 180° pulse length [μs] | 9.00 |
| $^{1}H$ LG decoupling power [kHz] | 85 |
| $^{1}H$ LG offset [kHz] | 60.1 |
| Scans | 32 |
| Experimental time ($S_0$ and S) [hours] | ~38.5 |
| Analysis software | SPARKY 3.134 |

INADEQUATE-REDOR experimental parameters

| | |
|---|---|
| Dephasing times [μs] | 429, 571, 714, 1143, 1286, 1429, 1571, 1714 |
| Acquisition mode in indirect dimension | States |
| Acquisition points (t1/t2) | 450/3988 |
| Acquisition time (t1/t2) [ms] | 3.75/20.00 |
| Spectral Width F1/F2 [kHz] | 60/100 |
| τ† [ms] | 3.50 |
| z-filter duration [μs] | 71.43 |
| Scans | 32 |
| Experimental time ($S_0$ and S) [hours] | ~24 |
| Analysis software | POKY[2] |

† See Figure S1

The noise level was estimated by the noise calculation function of SPARKY. The noise was calculated as the average of 10 estimations.

The phase scheme of the dephasing pulses of the REDOR block was XY8.[3]

Complete phase cycles are shown explicitly in Figure S1.

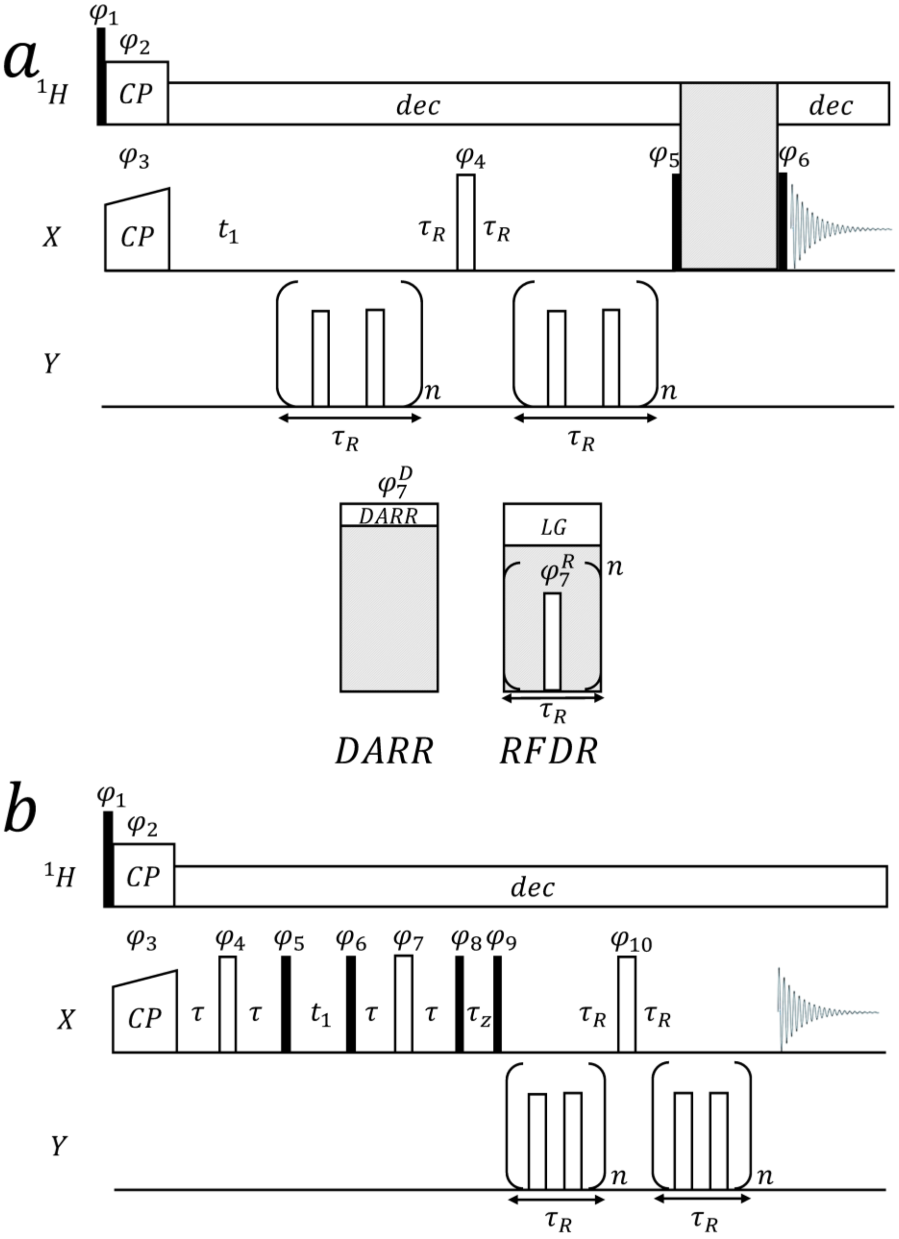

**Figure S1**. Pulse sequences for the measurement of C-N dipolar coupling constants. Black and white rectangles represent 90° and 180° pulses, respectively. CP stands for Cross Polarisation block, and 'dec' stands for $^1$H decoupling. $\tau_R$ is the MAS rotor period. (a) X{Y}-X REDOR-DARR and REDOR-RFDR pulse sequences. The two applied schemes are identical with the exception of the mixing block represented by the grey rectangle. For DARR mixing the $^1$H radio-frequency irradiation strength $\nu_1$ is applied during mixing at the rotary resonance condition $\nu_1 = \tau_R^{-1}$. For RFDR, rotor-synchronous 180° pulses are applied on the X channel ($^{13}$C in our experiments) and a homonuclear $^1$H Lee-Goldburg decoupling is employed. $\tau_R$ is the MAS rotor period. $\varphi_1 = \{90°, 270°\}, \varphi_2 = \varphi_7^D = \{0°\}, \varphi_3 = \{90°\}, \varphi_4 = \{90°\}_2\ \{180°\}_2, \varphi_5 = \{0°\}_4\ \{180°\}_4, \varphi_6 = \{90°\}$. For the RFDR pulses, $\varphi_7^R = XY - 8$ was employed. (b) X-X{Y} INADEQUATE-REDOR: $\varphi_1 = \{90°, 270°\},\ \varphi_2 = \varphi_6 = \varphi_8 = \varphi_9 = \{0°\}, \varphi_3 = \{90°\}_2\ \{180°\}_2\ \{270°\}_2\ \{0°\}_2, \varphi_4 = \varphi_5 = \{0°\}_2\ \{90°\}_2\ \{180°\}_2\ \{270°\}_2, \varphi_7 = \{90°\}_{16}\ \{270°\}_{16}, \varphi_{10} = \{90°\}$.

### S2 – Matlab scripts for fitting the REDOR curves

The Matlab script I is used to analyse the results from the 'Cross Peaks Analysis Tool' of RAVEN.[4] The input is a string array named 'results'. The required format for 'n' crosspeaks with 'k' dephasing times is

| $Crosspeak\ identifier$ | $\delta_1$ | $\delta_2$ | $S(\tau_1)$ | $S_0(\tau_1)$ | $S(\tau_2)$ | $S_0(\tau_2)$ | $\cdots$ | $S(\tau_k)$ | $S_0(\tau_k)$ |
|---|---|---|---|---|---|---|---|---|---|
| $Crosspeak\ \#1$ | $\delta_{1,1}$ | $\delta_{1,2}$ | $intensity$ | $intensity$ | $intensity$ | $intensity$ | $\cdots$ | $intensity$ | $intensity$ |
| $Crosspeak\ \#2$ | $\delta_{2,1}$ | $\delta_{2,2}$ | $intensity$ | $intensity$ | $intensity$ | $intensity$ | $\cdots$ | $intensity$ | $intensity$ |
| $\vdots$ | $\vdots$ | $\vdots$ | $intensity$ | $intensity$ | $intensity$ | $intensity$ | $\cdots$ | $intensity$ | $intensity$ |
| $Crosspeak\ \#n$ | $\delta_{n,1}$ | $\delta_{n,2}$ | $intensity$ | $intensity$ | $intensity$ | $intensity$ | $\cdots$ | $intensity$ | $intensity$ |
| | | | $noise$ | $noise$ | $noise$ | $noise$ | $\cdots$ | $noise$ | $noise$ |

The dephasing times $\tau_j$ are in units of μs.

An example of actual data:

| | | | | | | | | | | | | | | | | |
|---|---|---|---|---|---|---|---|---|---|---|---|---|---|---|---|---|
| | | | 428.6 | 428.6 | 571.4 | 571.4 | 857.1 | 857.1 | 1000 | 1000 | 1142.9 | 1142.9 | 1285.7 | 1285.7 | 1571.4 | 1571.4 |
| K8Ca-Cb | 59.81 | 32.79 | 13017212 | 14715122 | 11799954 | 14743988 | 9789653 | 13709820 | 8384717 | 15047111 | 6578601 | 13876720 | 5434314 | 13570522 | 3659389 | 12436655 |
| K8Ca-Cg | 59.93 | 25 | 3099990 | 3658567 | 2426522 | 3076466 | 2343983 | 3244493 | 1783701 | 3460326 | 1288482 | 3093055 | 1218291 | 2791504 | 376956.2 | 2825963 |
| A10Ca-Cb | 54.94 | 20.8 | 13652908 | 15166148 | 11295424 | 14656649 | 8781171 | 13244710 | 8132726 | 13724887 | 6385731 | 13374704 | 6363556 | 12789240 | 4084359 | 11535716 |
| | | | 204400 | 209300 | 198100 | 190500 | 192600 | 178700 | 206000 | 179400 | 206600 | 203200 | 192600 | 185500 | 182700 | 177500 |

The script calculates 1-$S_d$ for each data point, and then calculates the $\chi^2_{red}$ value for various dipolar coupling constants according to

$$\chi^2_{red}(d) = \frac{1}{N-1}\sum_{i}^{N}\frac{(O_i^2 - F_i^2)}{\sigma_i^2},$$

with 'd' being a specific calculated dipolar coupling constant, 'N' being the number of points in the REDOR dimension, 'O' being the observed value $1 - S_d(\tau_n)$, 'F' being the theoretical value of $1 - S_d(\tau_n)$ corresponding to the specific d according to the universal curve, and 'σ' is the error, calculated by

$$\sigma_i = |S_{d,i}|\sqrt{\left(\frac{n_{s,i}}{S_i}\right)^2 + \left(\frac{n_{s_0,i}}{S_{0,i}}\right)^2},$$

with $n_{s,i}$ and $n_{s_0,i}$ being the estimated noise level of the 'S' and '$S_0$' spectra respectively.

The effective dipolar coupling constant of a specific spin, $d$, is the one corresponding to the minimum $\chi^2_{red}$ value ($\chi^2_{red,min}$). The error of the fit was calculated according to the span of dipolar coupling constants that fulfil $\chi^2_{red}(d) = 2 \cdot \chi^2_{red,min}(d_{eff})$.

**Matlab script I**:

```
a = size(results);

%Creating a matrix of dephasing times in units of sec
j = 4;
deph = [];
data = ["crosspeak"];
while j<a(2)
```

```
    deph = [deph, results(1,j)];
    data = [data, append(results(1,j)," µs")];
    j = j+2;
end
deph = 10^-6*str2double(deph);

%Set the screened dipolar constants
res = 1; %The resolution in the screened dipolar constants in Hz
dip = [5:res:2000]; %array of wanted dipolar constants range in Hz
tau = [0:0.0010:max(deph)*10^3]; %time in ms

%Creating a string array of crosspeaks, sd and error matrix
m = 2;
error = [];
while m<a(1)
        temp = results(m,1);
        line = [temp];
        linen = [];
        j = 4;
        while j<a(2)
            s = str2double(results(m,j));
            n = str2double(results(a(1),j));
            s0 = str2double(results(m,j+1));
            n0 = str2double(results(a(1),j+1));
            sd = 1-s/s0;
            noise = abs(s/s0)*sqrt((n0/s0)^2+(n/s)^2);
            line = [line, sd];
            linen = [linen,noise];
            j = j+2;
        end
        data = [data; line];
        error = [error;linen];
        m = m+1;
end

%Calculate χ^2_red for each crosspeak and determine the dipole constant
coef = 0.25*sqrt(2)*pi();
[b,~] = size(data);
[~,c] = size(deph);
[~,d] = size(dip);
i = 1;
while i<d+1
    column = [(dip(i))];
    m = 2;
    while m<b+1
        j = 2;
        r = 0;
        while j<c+2
            sd = str2double(data(m,j));
            lam = sqrt(2)*deph(j-1)*dip(i);
            SD = 1-coef*besselj(0.25,lam)*besselj(-0.25,lam);
            temperr = error(m-1,j-1);
            r = r + ((SD-sd)/temperr)^2;
            j = j+1;
        end
```

```
        r = r/(c-1);
        column = [column;r];
        m = m+1;
    end
    data = [data,column];
    i = i+1;
end

%Calculate the error in the effective dipolar interaction
RMSDtable = str2double(data(:,(c+2):(c+d+1)));
m = 2;
column1 = ["Dipole constant in Hz"];
column2 = ["χ^2_red"];
column3 = ["Error"];

while m<b+1
    line = RMSDtable(m,:);
    [M,I] = min(line);

    err = 2*M;
    p = I;
    while line(p)<=err
        p = p-1;
        if p==0
            leftedge = dip(1);
            break
        end
        leftedge = dip(p+1);
    end

    p = I;
    while line(p)<=err
        p = p+1;
        if p>d
            rightedge = dip(1);
            break
        end
        rightedge = dip(p-1);
    end

    if (rightedge - dip(I)) == 0
        plus = append("^{+",num2str(res),"}");
    else
        plus = append("^{+",num2str(rightedge - dip(I)),"}");
    end
    if (dip(I) - leftedge) == 0
        minus = append("_{-",num2str(res),"}");
    else
        minus = append("_{-",num2str(dip(I) - leftedge),"}");
    end
    column3 = [column3;append(plus,minus)];

    M = num2str(M);
    I = num2str(dip(I));
    column1 = [column1; append(I)];
```

```
    column2 = [column2; M];
    m = m+1;
end
data = [data, column1, column2, column3];

%Plot the curve with the data points on it for each crosspeak
    f1 = figure;
    figure(f1);
    t=tiledlayout('flow');
    nexttile
    m = 2;
    while m<b+1
        dipolar = str2num(data(m,c+d+2));
        lam = sqrt(2)*dipolar.*tau*10^-3;
        sd = 1-coef*besselj(0.25,lam).*besselj(-0.25,lam);
        plot(tau,sd,'LineWidth', 1.5);
        x = deph*10^3;
        hold on
        j = 2;
        y = [];
        while j<(c+2)
            y = [y,str2double(data(m,j))];
            j = j+1;
        end
        temperr = error(m-1,:);
        errorbar(x,y,temperr, temperr,'o');
        set(gca,'FontSize',15);
        dipolar = num2str(dipolar);
        [~,f] = size(data);
        s = append(data(m,1),'; ', dipolar, data(m,f), ' ', 'Hz');
        title(s);
        xlabel("dephasing time [ms]", 'FontSize', 15);
        ylabel("1-S/S_0", 'FontSize', 15);
        if m==b(1)
            hold off
        else
            nexttile
        end
        m = m+1;
    end

%Plot χ^2_red(d) for each crosspeak
f2 = figure;
figure(f2);
t2 = tiledlayout('flow');
nexttile
m = 2;
while m<b+1
    i = c+2;
    y = [];
    while i<(d+c+2)
        y = [y, str2num(data(m,i))];
        i = i+1;
    end
    scatter(dip,y, 'LineWidth', 1.5);
```

```
    title(data(m,1));
    xlabel("Dipolar constant in Hz", 'FontSize', 15);
    ylabel("χ^2_r_e_d", 'FontSize', 15);
    if m==b(1)
        hold off
    else
        nexttile
    end
    m = m+1;
end
```

**Matlab script II**: This script is used in order to convert the 'matches' output file of RAVEN into a new list in the required format for the 'Cross Peaks Analysis Tool' of RAVEN.

```
%Takes 'matches' file from Raven and turns it into a list of unambiguous crosspeaks
as 'new.list', that can be used for the crosspeak analysis tool of Raven.

A = readlines('matches.txt');
k = size(A);
i = 2;
j = 2;
c = [];
while i<k(1)
    temp = [];
    if A(i) ~= ""
        j = i;
        while A(j) ~= ""
            temp = [temp; A(j)];
            j = j+1;
        end
    end
    [token1, remain] = strtok(temp);
    [token, remain] = strtok(remain);
    if all(token==token(1))
        c = [c;token];
    end
    i=j+1;
end

t = size(c);
i = 1;
d = ["Assignment" "w1" "w2"];
while i<t(1)+1
    [name1,remain] = strtok(c(i),'(');
    [cs1,remain] = strtok(remain,'-');
    [name2,cs2] = strtok(remain,'(');
    name = append(name1,name2);
    w1 = erase(cs1,'(');
    w1 = erase(w1,')');
    w2 = erase(cs2,'(');
    w2 = erase(w2,')');
    i = i+1;
```

```
    d = [d;name, w1, w2];
end

fileID = fopen('new.list', 'w');
fprintf(fileID, '%s \n', 'CC');
fprintf(fileID,'%s %s %s \n',d');
fclose(fileID);
```

Both scripts were written on version R2021b of Matlab by MathWorks Inc.

### S3 – DQ extension for RAVEN and POKY

Both RAVEN and POKY do not support DQ data. In order to make the analysis process for the INADEQUATE-REDOR p3D experiment automated (or in general use RAVEN or POKY for DQ data), we wrote a Python script that manipulates the DQ data to fit the necessary formats for the two programmes.

**DQ data in RAVEN:** The script creates a new sequence file with additional residues that allows the representation of DQ data by duplicating the original sequence. Since RAVEN does not support assignment tables with resonances that do not correspond to predefined conventional atoms as they appear in the BMBR, the modified table labels all possible combinations of bonded atoms, which can potentially be obtained from J-based DQ correlations, as single spins. This is demonstrated in Figure S3.1.

| | | $C$ | $C_\alpha$ |
|---|---|---|---|
| Original | $M1$ | $a_0$ | $a_1$ |
| | $A2$ | $b_0$ | $b_1$ |
| | $S3$ | $c_0$ | $c_1$ |
| | $N4$ | $d_0$ | $d_1$ |
| | $M5$ | $e_0$ | $e_1$ |
| | $R6$ | $f_0$ | $f_1$ |
| DQ | &$M7$ | $a_0 + a_1$ | $a_1 + a_2$ |
| | &$A8$ | $b_0 + b_1$ | $b_1 + b_2$ |
| | &$S9$ | $c_0 + c_1$ | $c_1 + c_2$ |
| | &$N10$ | $d_0 + d_1$ | $d_1 + d_2$ |
| | &$M11$ | $e_0 + e_1$ | $e_1 + e_2$ |
| | &$R12$ | $f_0 + f_1$ | $f_1 + f_2$ |

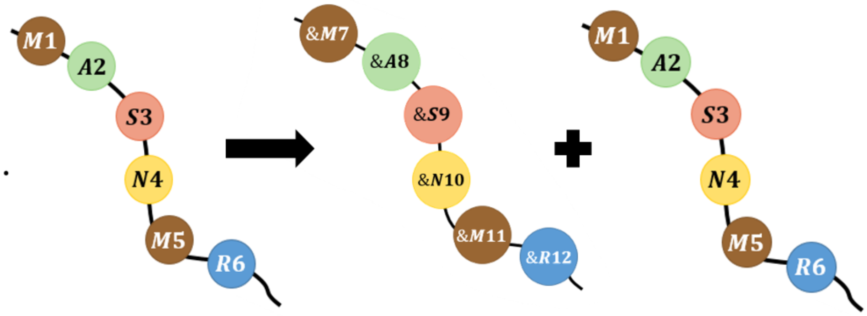

Figure S3.1. Illustration of the conversion process on a six-residue-long sequence. The left panel demonstrates the sequence duplication. The script denotes the fictitious residues with '&' and calculates the DQ resonances based on the original assignment table. The subscript '2' in $a_2$, $b_2$, etc. corresponds to $C_\beta$. Additional $a_k$, $b_k$, etc. are generated according to the length of the particular amino acids.

The automated assignment process done by RAVEN yields a 'matches' file that includes all possible assignment options for resonances with the addition of the DQ data. The script then converts the fictitious labels ("new" amino acids and resonances) into the proper labels. In the process it filters out assignment options that are either impossible in J-based INADEQUATE (inter-residues contacts) or improbable ($^2J_{CC}$ and higher contacts). The user can also specify the $^{13}C$ isotopic labelling scheme that was used for the sample preparation, including probabilities resulting from scrambling. The script filters out improbable assignment options according to a scrambling threshold (joint probability for an assigned spin-pair) selected by the user. The result is a filtered 'match' file with DQ data. This process is illustrated in Figure S3.2.

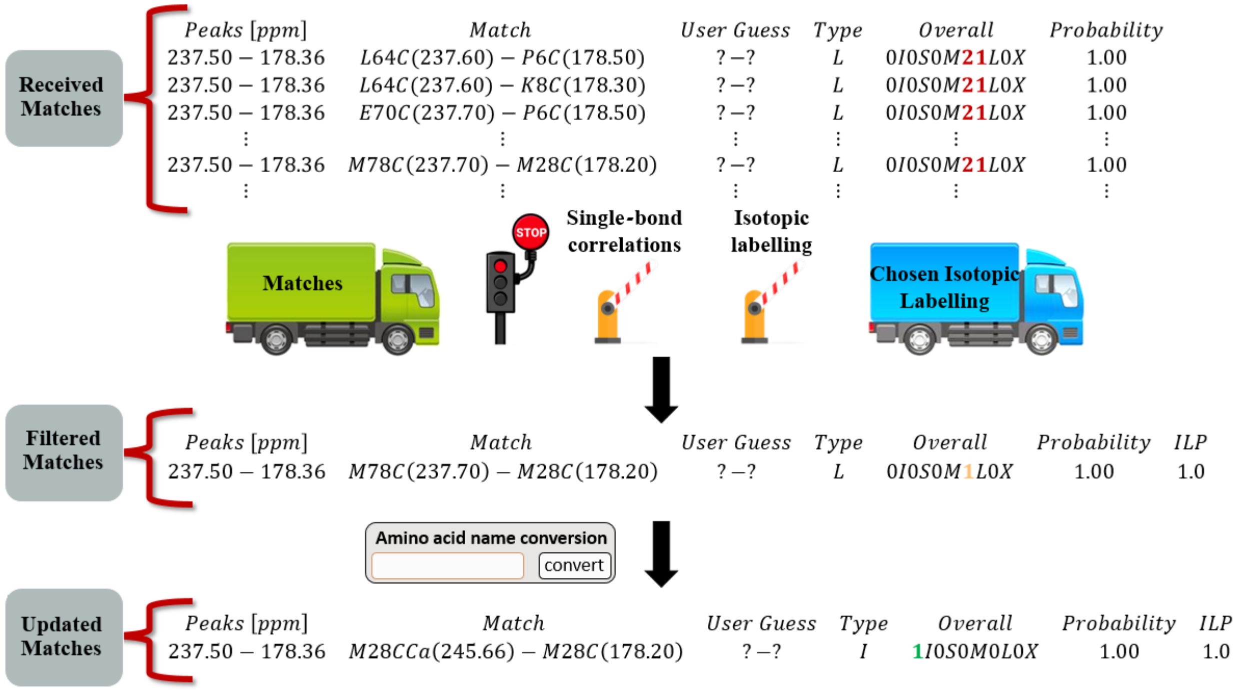

**Figure S3.2**. Illustration of the conversion of fictitious-DQ notation of the RAVEN 'matches' file into an explicit DQ peak list. The First block contains all possible assignment options for crosspeaks in the 50-residues long coat protein from the fd-Y21M phage coat protein. Matches are filtered by eliminating the improbable assignments based on the labelling scheme and on chemical bonds. Then, the labels of the amino acids are converted into their proper labels. Here M28CCa(245.66) represents a double-quantum peak correlation of methionine #28 with the sum of C (carbonyl) and Cα in the indirect dimension at 245.66 ppm, and the carbonyl resonance at 178.2 ppm.

The python script supports two additional RAVEN features that are non-specific to the DQ data. (i) It can convert 'matches' file into the 'Cross Peak Analysis Tool' format of RAVEN, and (ii) it can compare the 'matches' file with the peak list to identify the crosspeaks that were not assigned by RAVEN.

**DQ data in POKY:** Another function in the script creates a POKY-formatted assignment table with DQ resonances. The result can be used to analyse INADEQUATE spectra in POKY, as demonstrated in Figure S3.3.

An independent script suitable for POKY alone has also been written and is available in the GitHub repository.

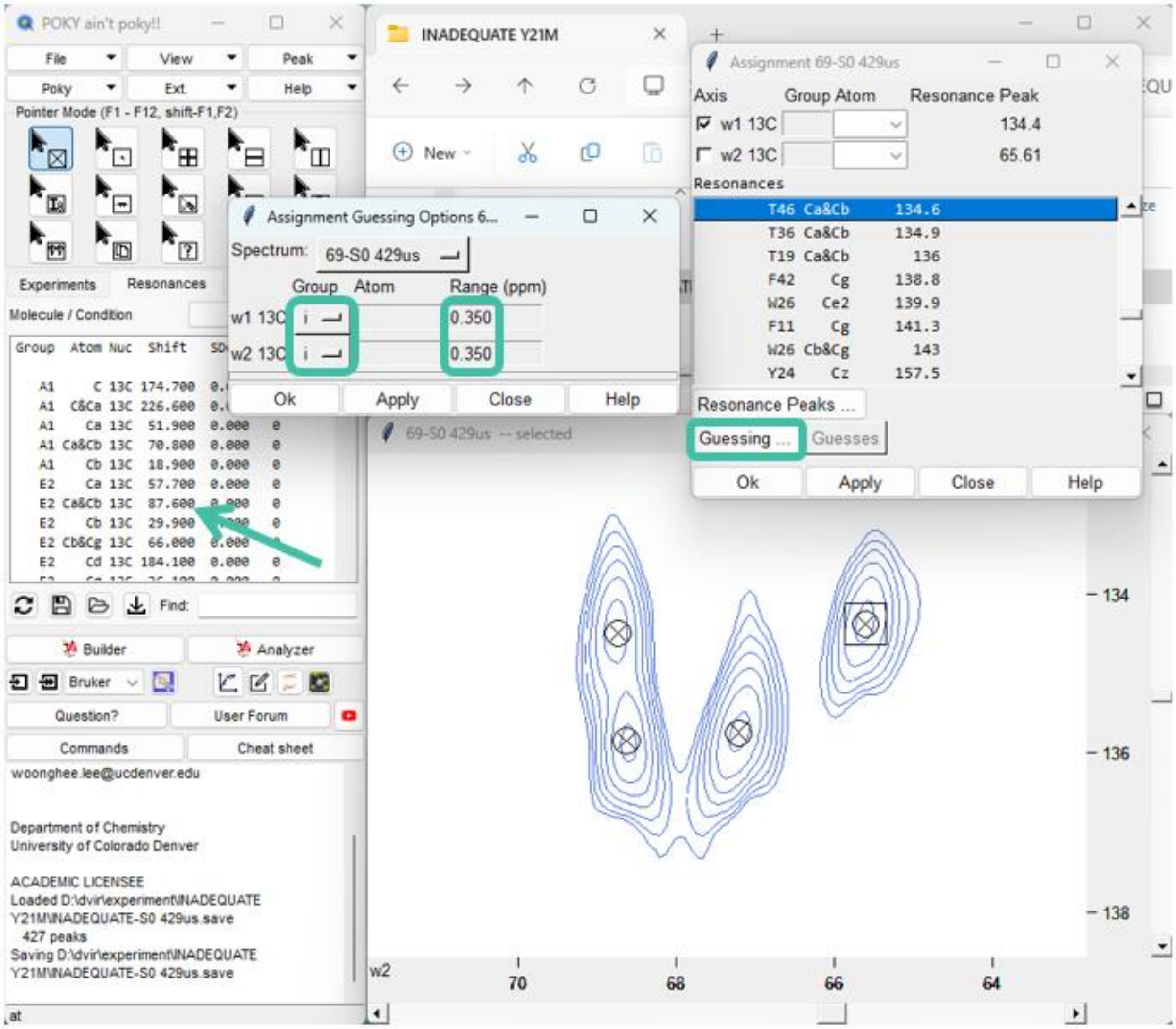

**Figure S3.3**. Screenshots of the POKY software interface. The spectrum is from the '$S_0$' INADEQUATE-REDOR experiment collected with a REDOR dephasing time of 429μs, conducted on the intact fd-Y21M phage. The arrow points at the modified assignment table that supports the DQ data. The green rectangles show the function in POKY that guesses the assignment of crosspeaks according to the modified assignment table.

The full script, along with tutorial videos and a manual can be found in GitHub, https://github.com/amirgoldbourt/DQ-extension-for-RAVEN-and-POKY.

### S4 – Validity of linear prediction and the universal REDOR curve approach in the analysis of p3D experiments

**Validity of linear prediction:** Linear prediction is used in order to increase the resolution in the indirect dimension by reducing the linewidths thereby improving sensitivity.[5] The linear prediction was needed for the INADEQUATE and RFDR based experiments, and it allowed the analysis of more residues. In order to ensure that the linear prediction does not change the results (quantification of the REDOR curve), we calculated 1-$S_d$ for crosspeaks that were non-ambiguous before the linear prediction and compared the results to those obtained from a spectrum with linear prediction.

Figure S4.1 shows the build-up curves for three crosspeaks with and without linear prediction as an example. The results of the fit are summarised in table S4 and show that the difference is smaller than the experimental errors, thus validating the usage of linear prediction.

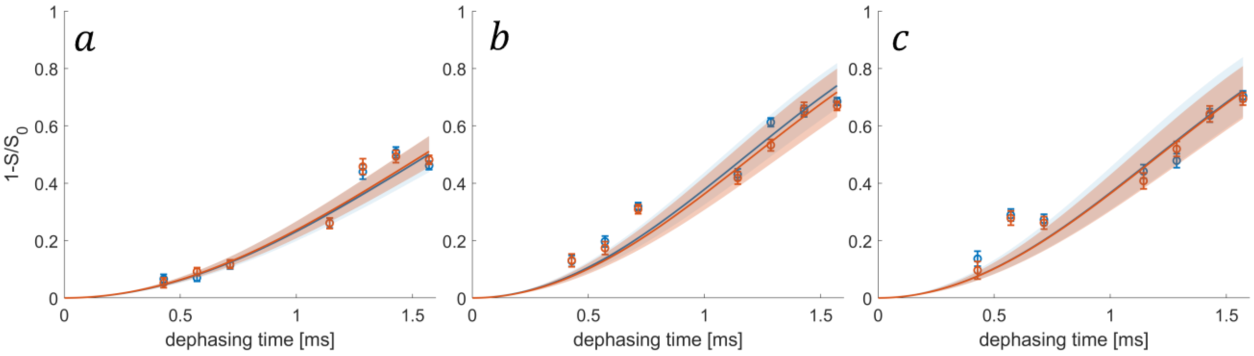

**Figure S4.1**. INADEQUTE-REDOR build up curves of three crosspeaks: (a) Q15 Cγ&Cδ-Cδ, (b) T46 Cα&Cβ-Cα, (c) T19 Cα&Cβ-Cα with linear prediction (blue) and without linear prediction (orange). The experimental data is shown in circles along with the errors. The curves represent the best fit to the universal REDOR curve. The shaded areas correspond to the level of confidence derived by the errors summarised in table S4.1.

| Crosspeak | '$d_{eff}$' with linear prediction | '$d_{eff}$' without linear prediction |
|---|---|---|
| Q15 Cγ&Cδ-Cδ | $488^{+39}_{-39}$ | $494^{+34}_{-34}$ |
| T46 Cα&Cβ-Cα | $645^{+59}_{-54}$ | $629^{+60}_{-58}$ |
| T19 Cα&Cβ-Cα | $633^{+77}_{-72}$ | $630^{+66}_{-63}$ |

**Table S4.1**. Effective dipolar coupling constants 'd' calculated by fits to the universal REDOR curve (in units of Hz) for three different crosspeaks in INADEQUATE-REDOR experiments with and without linear prediction.

**Validity of the universal REDOR curve**: The fit to the universal REDOR curves assumes an isolated two-spin system. In order to validate this assumption, we used the NMR simulation package SIMPSON[6] and its original REDOR script. First, we simulated a three-spin system consisting of one $^{13}C$ spin and two $^{15}N$ spins ($CN_2$). The strength of the heteronuclear dipolar coupling constants were chosen to be 1005Hz and 200Hz, derived from the typical distances between Cα and the two N spins of the same the adjacent amino acid. Figure S4.2a shows the simulated build-up curve along the best fit. We also simulated a four-spin system with three $^{13}C$ spins (representing C, Cα and Cβ) and one $^{15}N$ ($C_3N$). The homonuclear dipolar interactions Cα-C and Cα-Cβ were chosen to be 2250Hz (corresponding to a distance of 1.5Å), and the heteronuclear Cα-N dipolar interaction was set to 1005Hz. Figure S4.2b shows the simulated build-up curve with the fit. In both simulations an error of 5% was added to the data points in order to resemble experimental conditions.

For the $CN_2$ system the best fit yielded a result of $d_{eff} = 1007Hz$, a deviation of 2Hz. The $C_3N$ system yielded an effective dipolar coupling constant of $d_{eff} = 1099Hz$, a deviation of 4Hz. These deviations are both smaller than the experimental errors, thus validating the assumptions.

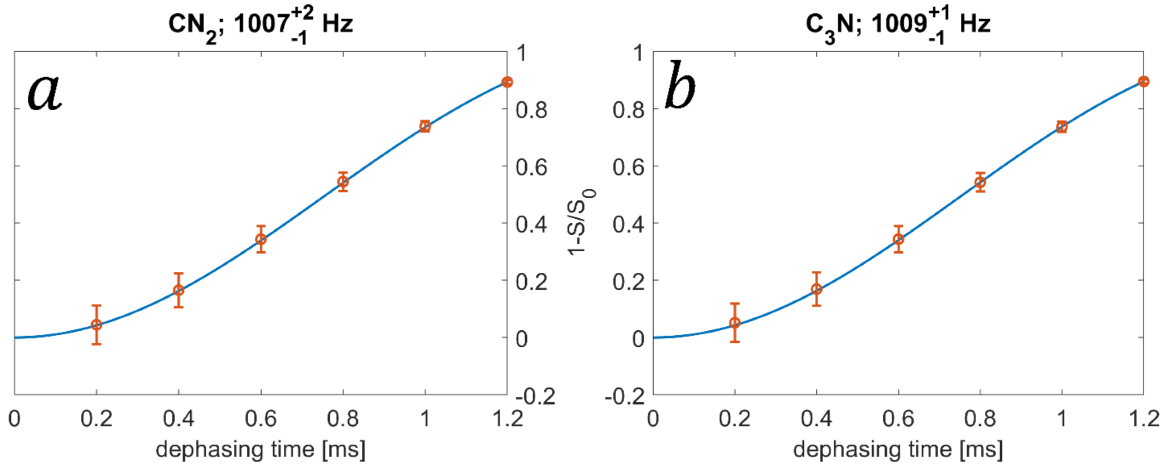

**Figure S4.2**. Build up curves of simulated systems (orange) along with the best fit to the universal REDOR curve (blue). (a) A system of Cα with two $^{15}N$ spins. The dipolar coupling constants were 1005Hz and 200Hz, corresponding to typical values in the backbone of proteins. (b) A system of C, Cα and Cβ spins along with a single $^{15}N$ spin attached to the Cα. The homonuclear and heteronuclear dipolar coupling constants were 2250Hz and 1005Hz, respectively.

### S5 – Detailed example of the data analysis procedure

The general steps for data analysis are as follows:

- The spectra are recorded, Fourier transformed, and the required crosspeaks are peak-picked and recorded. The crosspeaks are then assigned automatically by RAVEN according to the assignment table.
- The RAVEN 'matches' file is converted into the required format (using the Matlab script I in S2) for the 'Cross Peak Analysis Tool' of RAVEN.
- The assigned peak list is uploaded onto the 'Cross Peak Analysis Tool' of RAVEN along with all the UCSF files (the format is generated by the nmrPipe 'ucsfdata' tool) corresponding to the p3D experiments.
- The build-up curves are saved and uploaded into Matlab (using script II) as a string array that includes the noise level of each experiment.
- The build-up curves are created along with a $\chi^2_{red}$ plot.

Figure S5 illustrates the stages of data analysis on the P6Cδ-Cγ crosspeak detected in the REDOR-DARR experiments.

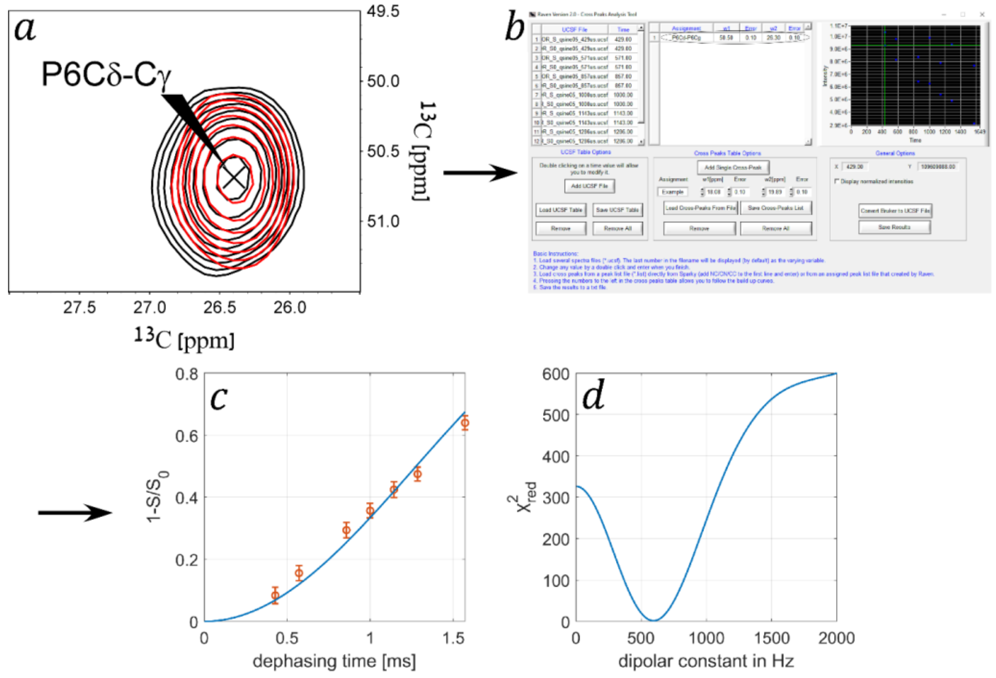

**Figure S5**. An illustration of the data analysis process. (a) The crosspeak P6Cδ-Cγ is identified in the DARR-REDOR spectra. The black and RED are the '$S_0$' and 'S' spectra respectively acquired with a REDOR dephasing time of 857μs. Contour levels are drawn at multiples of 1.2 with the lowest matching an SNR of 6. (b) A screenshot of RAVEN Cross Peak Analysis Tool. For clarity, only the data of the crosspeak in (a) appear in the panel. (c) A REDOR build-up curve created by calculating 1-$S_d$ using the results from (b) and the Matlab script I. The data points (1-$S_d$) are shown in orange with the errors, and the best fit curve is shown in blue. The fit yielded $d_{eff} = 595^{+28}_{-27}$. (d) $\chi^2_{red}$ plot for all screened dipolar coupling constants. Here $\chi^2_{red,min}(d_{eff} = 595Hz) = 1.66$.

The best fit yielded a result of $d_{eff} = 595^{+28}_{-27}$, which corresponds to an order parameter of $0.59^{+0.03}_{-0.03}$. In total, three crosspeaks involving the dephasing of P6Cδ were recorded – one in each mixing scheme. After analysing all three crosspeaks, the average order parameter for the Cδ-N bond was measured to be $0.58^{+0.03}_{-0.03}$.

### S6 – Full results of dephasing curves

**Table S6.1**: Backbone Cα-N (and Cδ-N in P6) order parameters for the fd-Y21M phage coat protein. A dash represents a residue without data. The rigid limit is $d = 1005Hz$. The total amount of observed crosspeaks is indicated, along with 'D', 'R' and 'I' standing for the mixing scheme in which they appeared (D=DARR, R=RFDR, I=INADEQUATE) and the number of crosspeaks in each of the experiments (e.g. 3R: 3 peaks observed in REDOR-RFDR). Each pair of A7 and A35, D12 and L14, and I32 and I39 also has an addition of two ambiguous crosspeaks in both DARR and RFDR mixings that were analysed.

| Residue | Information on resolved crosspeaks | Order parameter | Amplitude (Motion on a cone) (°) | Amplitude (Motion in a cone) (°) |
|---|---|---|---|---|
| A1 | 1 (R) | $0.16^{+0.02}_{-0.03}$ | $48.4^{+1.2}_{-1.4}$ | $75.2^{+2.0}_{-2.2}$ |

| | | | | |
|---|---|---|---|---|
| D2 | 1 (R) | $0.22^{+0.04}_{-0.06}$ | $46.2^{+1.7}_{-2.1}$ | $70.8^{+3.3}_{-4.1}$ |
| G3 | 2 (R,I) | $0.26^{+0.04}_{-0.06}$ | $44.4^{+1.7}_{-2.2}$ | $67.5^{+3.1}_{-4.0}$ |
| D4 | 2 (R) | $0.36^{+0.02}_{-0.05}$ | $40.6^{+0.9}_{-1.0}$ | $60.7^{+1.6}_{-1.7}$ |
| D5 | 6 (D, 3R, 2I) | $0.50^{+0.03}_{-0.03}$ | $35.2^{+1.2}_{-1.3}$ | $51.7^{+2.0}_{-2.1}$ |
| P6 Cα | 5 (D, 3R, I) | $0.64^{+0.04}_{-0.04}$ | $29.5^{+1.7}_{-1.6}$ | $42.7^{+2.6}_{-2.4}$ |
| P6 Cδ | 3 (D, R, I) | $0.58^{+0.03}_{-0.03}$ | $31.8^{+0.5}_{-0.4}$ | $46.3^{+1.1}_{-1.1}$ |
| A7 | 3 (D, R, I) | $0.67^{+0.03}_{-0.03}$ | $27.9^{+0.9}_{-0.9}$ | $40.3^{+2.1}_{-2.2}$ |
| K8 | 7 (3D, 3R, I) | $0.69^{+0.02}_{-0.02}$ | $26.8^{+1.0}_{-0.9}$ | $38.7^{+1.6}_{-1.4}$ |
| A9 | | - | | |
| A10 | 4 (D, 2R, I) | $0.66^{+0.03}_{-0.03}$ | $28.3^{+1.4}_{-1.4}$ | $41.0^{+2.2}_{-2.1}$ |
| F11 | 5 (2D, 2R, I) | $0.66^{+0.05}_{-0.04}$ | $28.2^{+2.2}_{-1.9}$ | $40.8^{+3.3}_{-2.8}$ |
| D12 | 4 (2D, R, I) | $0.64^{+0.04}_{-0.03}$ | $29.4^{+1.1}_{-1.1}$ | $42.7^{+2.5}_{-2.4}$ |
| S13 | | - | | |
| L14 | 1 (I) | $0.64^{+0.04}_{-0.04}$ | $29.2^{+1.2}_{-1.1}$ | $42.3^{+2.6}_{-2.5}$ |
| Q15 | 3 (2D, I) | $0.65^{+0.03}_{-0.03}$ | $29.0^{+1.6}_{-1.5}$ | $42.1^{+2.4}_{-2.3}$ |
| A16 | 1 (D) | $0.66^{+0.05}_{-0.05}$ | $28.3^{+2.5}_{-2.4}$ | $41.0^{+3.8}_{-3.7}$ |
| S17 | | - | | |
| A18 | 1 (R) | $0.70^{+0.04}_{-0.04}$ | $26.4^{+2.0}_{-1.9}$ | $38.1^{+3.0}_{-2.9}$ |
| T19 | 5 (D, 2R, 2I) | $0.63^{+0.03}_{-0.03}$ | $29.9^{+1.2}_{-1.1}$ | $43.4^{+1.8}_{-1.8}$ |
| E20 | 4 (D, 2R, I) | $0.63^{+0.04}_{-0.04}$ | $29.6^{+1.9}_{-1.7}$ | $43.0^{+2.9}_{-2.6}$ |
| M21 | | - | | |
| I22 | 5 (2D, 2R, I) | $0.66^{+0.05}_{-0.04}$ | $28.3^{+2.1}_{-2.0}$ | $40.9^{+3.3}_{-3.0}$ |
| G23 | 2 (D, I) | $0.69^{+0.04}_{-0.04}$ | $27.1^{+2.1}_{-2.0}$ | $39.1^{+3.2}_{-3.0}$ |
| Y24 | | - | | |
| A25 | | - | | |
| W26 | 3 (D, R, I) | $0.65^{+0.06}_{-0.05}$ | $28.9^{+2.7}_{-2.4}$ | $41.9^{+4.1}_{-3.7}$ |
| A27 | | - | | |
| M28 | | - | | |
| V29 | 5 (2D, 2R, I) | $0.66^{+0.04}_{-0.04}$ | $28.6^{+1.9}_{-1.7}$ | $41.4^{+2.8}_{-2.6}$ |
| V30 | 5 (2D, R, 2I) | $0.64^{+0.14}_{-0.03}$ | $29.5^{+6.0}_{-1.3}$ | $42.8^{+9.2}_{-2.0}$ |
| V31 | 6 (2D, 3R, I) | $0.64^{+0.03}_{-0.03}$ | $29.3^{+1.4}_{-1.4}$ | $42.6^{+2.2}_{-2.1}$ |
| I32 | 3 (D, 2R) | $0.68^{+0.03}_{-0.03}$ | $27.7^{+1.0}_{-0.9}$ | $40.0^{+2.2}_{-2.0}$ |
| V33 | 1 (D) | $0.61^{+0.11}_{-0.09}$ | $30.7^{+4.6}_{-4.1}$ | $44.6^{+7.2}_{-6.4}$ |
| G34 | 2 (R, I) | $0.72^{+0.03}_{-0.03}$ | $25.8^{+1.7}_{-1.6}$ | $37.1^{+2.5}_{-2.4}$ |
| A35 | 4 (D, R, 2I) | $0.63^{+0.03}_{-0.03}$ | $29.7^{+0.9}_{-0.9}$ | $43.1^{+2.0}_{-1.9}$ |
| T36 | 2 (I) | $0.69^{+0.07}_{-0.06}$ | $27.2^{+3.2}_{-2.8}$ | $39.4^{+4.9}_{-4.2}$ |
| I37 | 3 (D, R, I) | $0.67^{+0.04}_{-0.04}$ | $27.9^{+1.7}_{-1.6}$ | $40.3^{+2.7}_{-2.5}$ |
| G38 | 3 (D, R, I) | $0.69^{+0.03}_{-0.03}$ | $27.0^{+1.6}_{-1.5}$ | $39.0^{+2.4}_{-2.3}$ |
| I39 | 3 (D, R, I) | $0.64^{+0.02}_{-0.02}$ | $29.3^{+0.8}_{-0.7}$ | $42.5^{+1.7}_{-1.6}$ |
| K40 | | - | | |
| L41 | 1 (R) | $0.65^{+0.04}_{-0.04}$ | $28.9^{+1.9}_{-1.8}$ | $41.8^{+3.0}_{-2.7}$ |
| F42 | 1 (D, R) | $0.70^{+0.07}_{-0.13}$ | $26.4^{+2.2}_{-4.1}$ | $38.1^{+5.2}_{-9.5}$ |
| K43 | | - | | |
| K44 | | - | | |
| F45 | | - | | |
| T46 | 2 (R) | $0.65^{+0.05}_{-0.05}$ | $29.0^{+2.4}_{-2.3}$ | $42.1^{+3.7}_{-3.6}$ |
| S47 | | - | | |
| K48 | | - | | |
| A49 | 3 (D, R, I) | $0.68^{+0.06}_{-0.05}$ | $27.7^{+2.7}_{-2.2}$ | $40.1^{+4.1}_{-3.3}$ |
| S50 | 4 (D, 2R, I) | $0.66^{+0.03}_{-0.03}$ | $28.5^{+1.4}_{-1.3}$ | $41.2^{+2.1}_{-1.9}$ |

**Table S6.2**. Sidechain order parameters discussed in the main text. The rigid limits for the dipolar coupling constants are 1332Hz for G, 1005 Hz for K, 1317 Hz for Q, 1116 Hz for W.

| #Residue | Amount of resolved crosspeaks | Bond | Order parameter | Amplitude (Motion on a cone) (°) | Amplitude (Motion in a cone) (°) |
|---|---|---|---|---|---|
| K8 | 2 (R, I) | Cε-Nε | $0.22^{+0.02}_{-0.02}$ | $46.3^{+0.8}_{-0.9}$ | $70.9^{+1.5}_{-1.8}$ |
| Q15 | 1 (I) | Cδ-Nε2 | $0.36^{+0.03}_{-0.03}$ | $40.9^{+1.3}_{-1.3}$ | $61.2^{+2.2}_{-2.2}$ |
| W26 | 2 (D, R) | Cε2-Nε1 | $0.75^{+0.10}_{-0.08}$ | $24.1^{+5.0}_{-4.1}$ | $34.7^{+7.4}_{-6.1}$ |
| K40 | 4 (2D, R, I) | Cε-Nζ | $0.57^{+0.03}_{-0.03}$ | $32.4^{+1.2}_{-1.2}$ | $47.3^{+1.9}_{-1.9}$ |
| K43 | 1 (R) | Cε-Nζ | $0.59^{+0.02}_{-0.04}$ | $31.6^{+1.0}_{-0.9}$ | $46.1^{+1.5}_{-1.4}$ |
| K44 | 2 (R, I) | Cε-Nζ | $0.47^{+0.04}_{-0.04}$ | $36.3^{+1.6}_{-1.7}$ | $53.5^{+2.7}_{-2.7}$ |

The following plots include all the fits to the dephasing curves of all crosspeaks we analysed in alphabetical order. The left panels in the three columns show the best-fit REDOR curve with the experimental data points, and the right panels show $\chi^2_{red}$. The crosspeak and the name of the mixing scheme are written in the title of the left panels, and the calculated effective dipolar interaction (and error) is written in the titles of the right panels. When ambiguous crosspeaks exist (not more than two options), both options are written in the title.

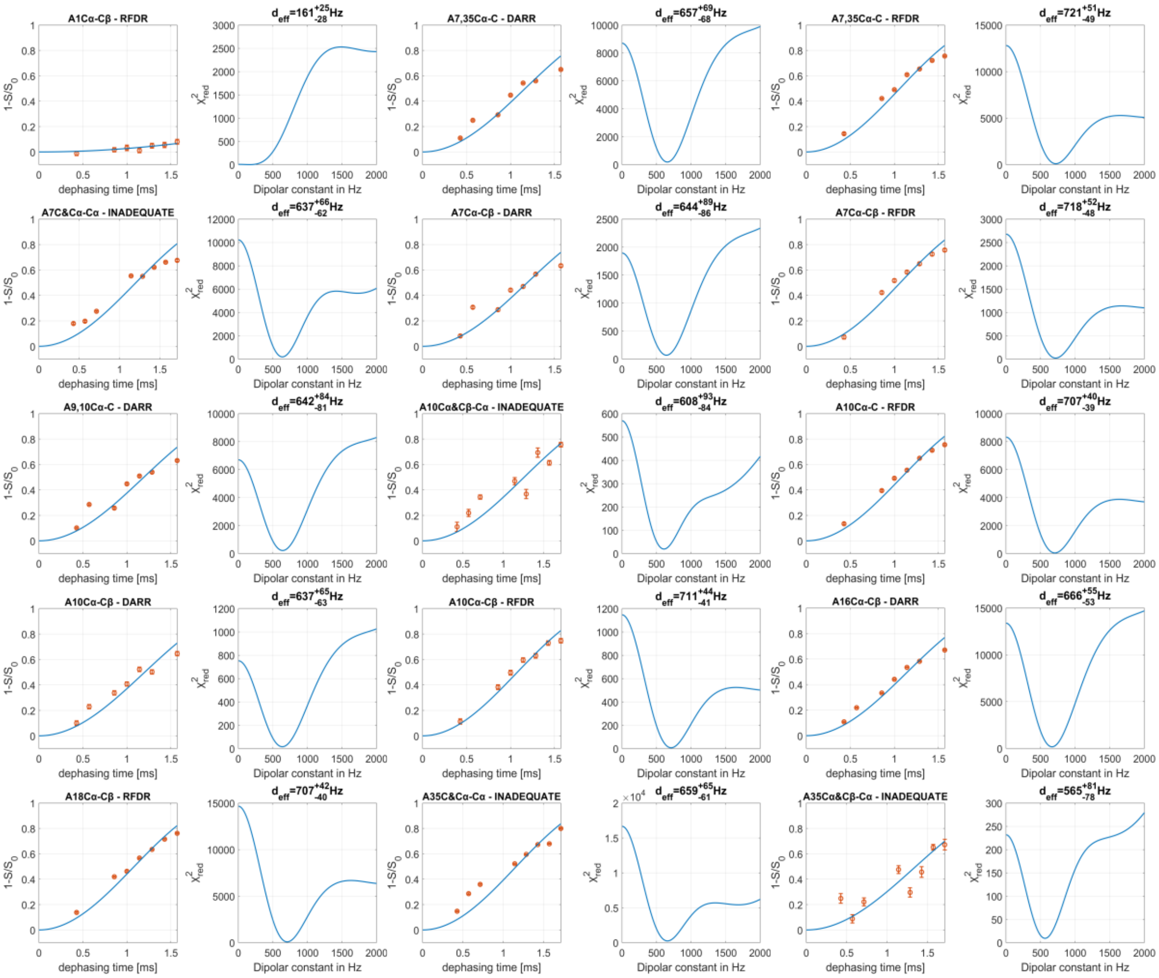

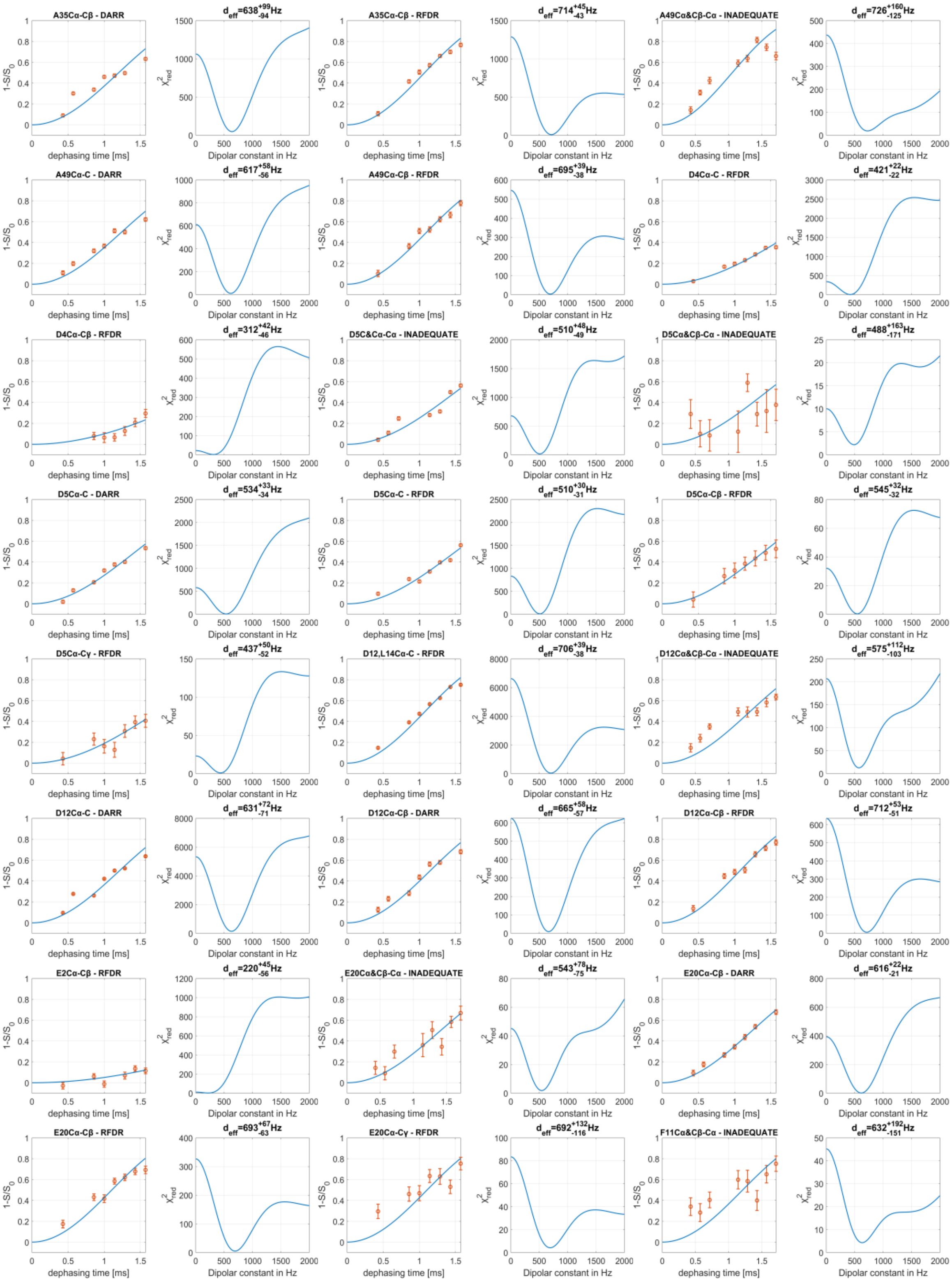
A35Cα-Cβ - DARR
$d_{eff}$=638$^{+99}_{-94}$Hz
A35Cα-Cβ - RFDR
$d_{eff}$=714$^{+45}_{-43}$Hz
A49Cα&Cβ-Cα - INADEQUATE
$d_{eff}$=726$^{+160}_{-125}$Hz
A49Cα-C - DARR
$d_{eff}$=617$^{+58}_{-56}$Hz
A49Cα-Cβ - RFDR
$d_{eff}$=695$^{+39}_{-38}$Hz
D4Cα-C - RFDR
$d_{eff}$=421$^{+22}_{-22}$Hz
D4Cα-Cβ - RFDR
$d_{eff}$=312$^{+42}_{-46}$Hz
D5C&Cα-Cα - INADEQUATE
$d_{eff}$=510$^{+48}_{-49}$Hz
D5Cα&Cβ-Cα - INADEQUATE
$d_{eff}$=488$^{+163}_{-171}$Hz
D5Cα-C - DARR
$d_{eff}$=534$^{+33}_{-34}$Hz
D5Cα-C - RFDR
$d_{eff}$=510$^{+30}_{-31}$Hz
D5Cα-Cβ - RFDR
$d_{eff}$=545$^{+32}_{-32}$Hz
D5Cα-Cγ - RFDR
$d_{eff}$=437$^{+50}_{-52}$Hz
D12,L14Cα-C - RFDR
$d_{eff}$=706$^{+39}_{-38}$Hz
D12Cα&Cβ-Cα - INADEQUATE
$d_{eff}$=575$^{+112}_{-103}$Hz
D12Cα-C - DARR
$d_{eff}$=631$^{+72}_{-71}$Hz
D12Cα-Cβ - DARR
$d_{eff}$=665$^{+58}_{-57}$Hz
D12Cα-Cβ - RFDR
$d_{eff}$=712$^{+53}_{-51}$Hz
E2Cα-Cβ - RFDR
$d_{eff}$=220$^{+45}_{-56}$Hz
E20Cα&Cβ-Cα - INADEQUATE
$d_{eff}$=543$^{+78}_{-75}$Hz
E20Cα-Cβ - DARR
$d_{eff}$=616$^{+22}_{-21}$Hz
E20Cα-Cβ - RFDR
$d_{eff}$=693$^{+67}_{-63}$Hz
E20Cα-Cγ - RFDR
$d_{eff}$=692$^{+132}_{-116}$Hz
F11Cα&Cβ-Cα - INADEQUATE
$d_{eff}$=632$^{+192}_{-151}$Hz
1-S/S$_0$
dephasing time [ms]
$\chi^2_{red}$
Dipolar constant in Hz

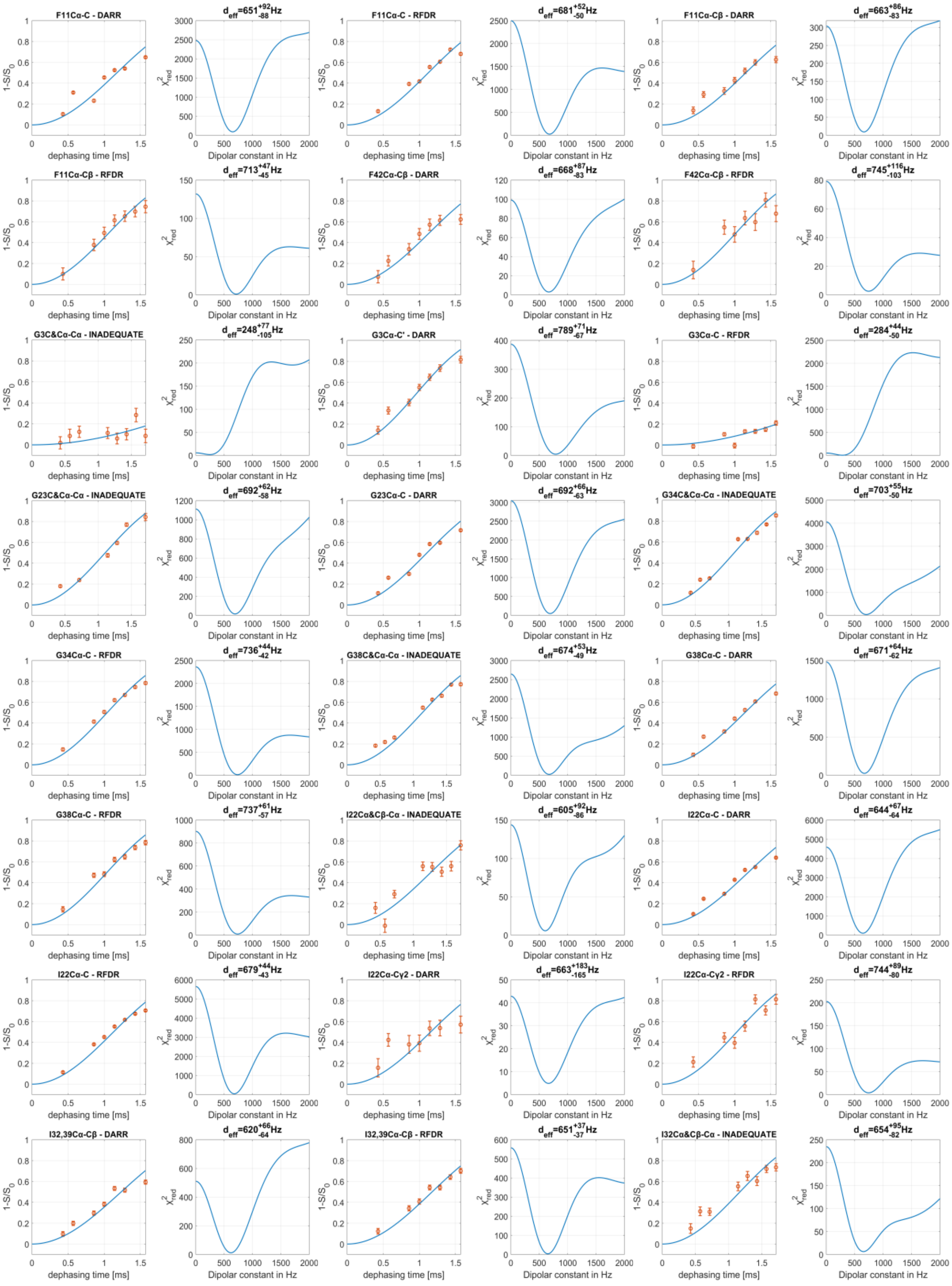
F11Cα-C - DARR
d_eff=651^{+92}_{-88}Hz
F11Cα-C - RFDR
d_eff=681^{+52}_{-50}Hz
F11Cα-Cβ - DARR
d_eff=663^{+86}_{-83}Hz
F11Cα-Cβ - RFDR
d_eff=713^{+47}_{-45}Hz
F42Cα-Cβ - DARR
d_eff=668^{+87}_{-83}Hz
F42Cα-Cβ - RFDR
d_eff=745^{+116}_{-103}Hz
G3C&Cα-Cα - INADEQUATE
d_eff=248^{+77}_{-105}Hz
G3Cα-C' - DARR
d_eff=789^{+71}_{-67}Hz
G3Cα-C - RFDR
d_eff=284^{+44}_{-50}Hz
G23C&Cα-Cα - INADEQUATE
d_eff=692^{+62}_{-58}Hz
G23Cα-C - DARR
d_eff=692^{+66}_{-63}Hz
G34C&Cα-Cα - INADEQUATE
d_eff=703^{+55}_{-50}Hz
G34Cα-C - RFDR
d_eff=736^{+44}_{-42}Hz
G38C&Cα-Cα - INADEQUATE
d_eff=674^{+53}_{-49}Hz
G38Cα-C - DARR
d_eff=671^{+64}_{-62}Hz
G38Cα-C - RFDR
d_eff=737^{+61}_{-57}Hz
I22Cα&Cβ-Cα - INADEQUATE
d_eff=605^{+92}_{-86}Hz
I22Cα-C - DARR
d_eff=644^{+67}_{-64}Hz
I22Cα-C - RFDR
d_eff=679^{+44}_{-43}Hz
I22Cα-Cγ2 - DARR
d_eff=663^{+183}_{-165}Hz
I22Cα-Cγ2 - RFDR
d_eff=744^{+89}_{-80}Hz
I32,39Cα-Cβ - DARR
d_eff=620^{+66}_{-64}Hz
I32,39Cα-Cβ - RFDR
d_eff=651^{+37}_{-37}Hz
I32Cα&Cβ-Cα - INADEQUATE
d_eff=654^{+95}_{-82}Hz
1-S/S0
dephasing time [ms]
χ²_red
Dipolar constant in Hz

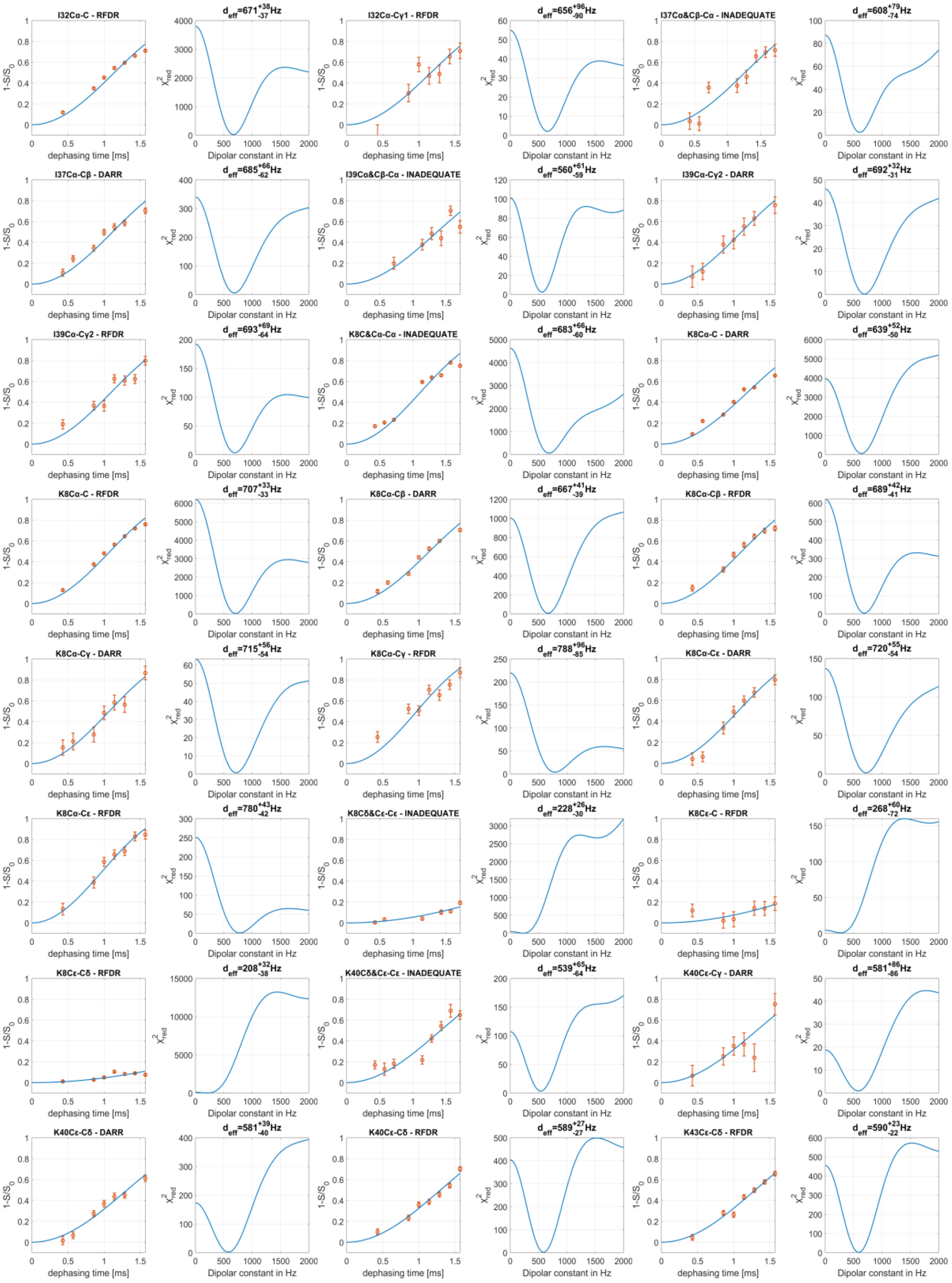
I32Cα-C - RFDR
$d_{eff}$=671$^{+38}_{-37}$Hz
I32Cα-Cγ1 - RFDR
$d_{eff}$=656$^{+96}_{-90}$Hz
I37Cα&Cβ-Cα - INADEQUATE
$d_{eff}$=608$^{+79}_{-74}$Hz
I37Cα-Cβ - DARR
$d_{eff}$=685$^{+66}_{-62}$Hz
I39Cα&Cβ-Cα - INADEQUATE
$d_{eff}$=560$^{+61}_{-59}$Hz
I39Cα-Cγ2 - DARR
$d_{eff}$=692$^{+32}_{-31}$Hz
I39Cα-Cγ2 - RFDR
$d_{eff}$=693$^{+69}_{-64}$Hz
K8C&Cα-Cα - INADEQUATE
$d_{eff}$=683$^{+66}_{-60}$Hz
K8Cα-C - DARR
$d_{eff}$=639$^{+52}_{-50}$Hz
K8Cα-C - RFDR
$d_{eff}$=707$^{+33}_{-33}$Hz
K8Cα-Cβ - DARR
$d_{eff}$=667$^{+41}_{-39}$Hz
K8Cα-Cβ - RFDR
$d_{eff}$=689$^{+42}_{-41}$Hz
K8Cα-Cγ - DARR
$d_{eff}$=715$^{+56}_{-54}$Hz
K8Cα-Cγ - RFDR
$d_{eff}$=788$^{+96}_{-85}$Hz
K8Cα-Cε - DARR
$d_{eff}$=720$^{+55}_{-54}$Hz
K8Cα-Cε - RFDR
$d_{eff}$=780$^{+43}_{-42}$Hz
K8Cδ&Cε-Cε - INADEQUATE
$d_{eff}$=228$^{+26}_{-30}$Hz
K8Cε-C - RFDR
$d_{eff}$=268$^{+60}_{-72}$Hz
K8Cε-Cδ - RFDR
$d_{eff}$=208$^{+32}_{-38}$Hz
K40Cδ&Cε-Cε - INADEQUATE
$d_{eff}$=539$^{+65}_{-64}$Hz
K40Cε-Cγ - DARR
$d_{eff}$=581$^{+86}_{-86}$Hz
K40Cε-Cδ - DARR
$d_{eff}$=581$^{+39}_{-40}$Hz
K40Cε-Cδ - RFDR
$d_{eff}$=589$^{+27}_{-27}$Hz
K43Cε-Cδ - RFDR
$d_{eff}$=590$^{+23}_{-22}$Hz
$1-S/S_0$
dephasing time [ms]
$\chi^2_{red}$
Dipolar constant in Hz

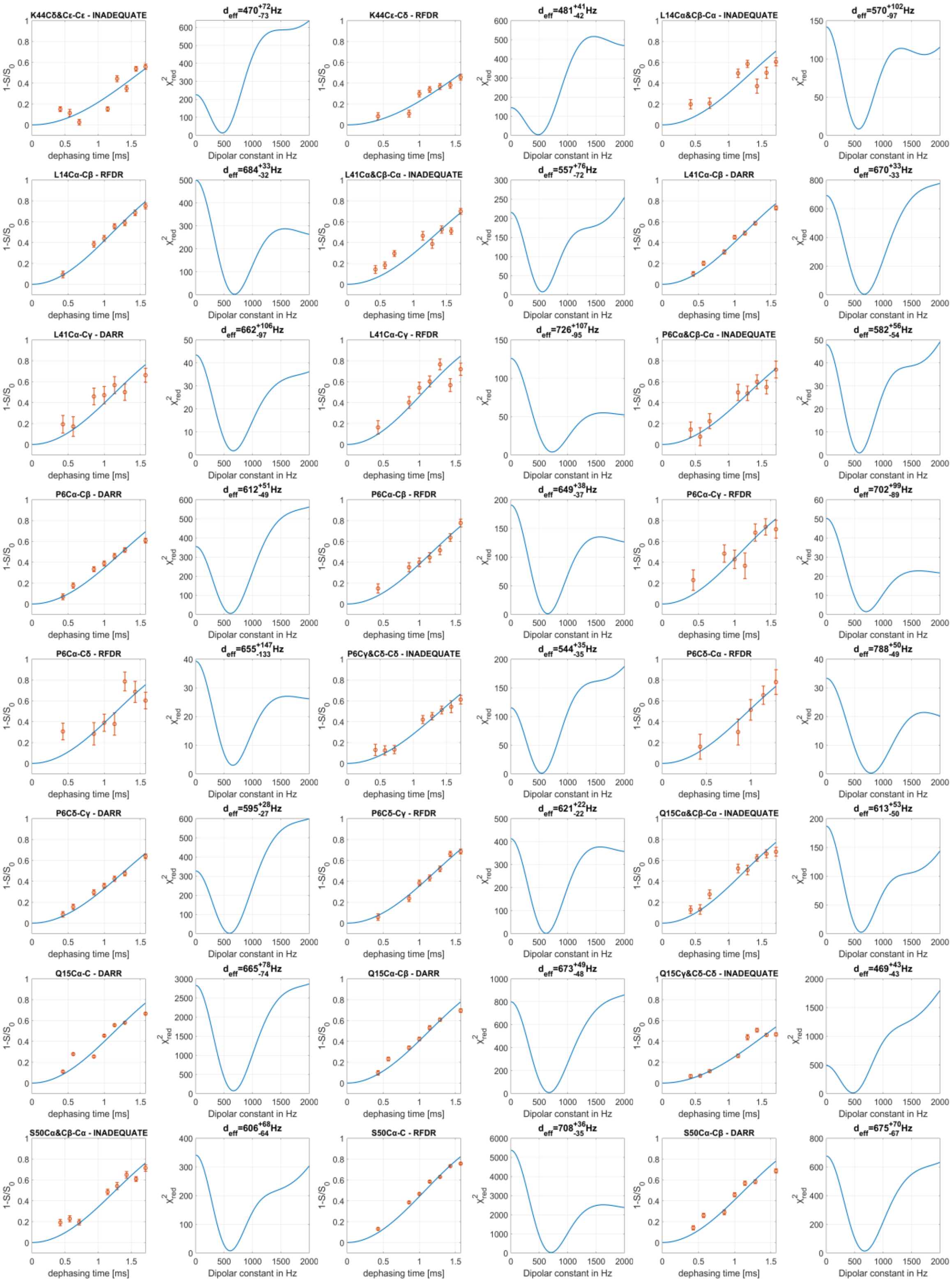
K44Cδ&Cε-Cε - INADEQUATE
$d_{eff}=470^{+72}_{-73}$Hz
K44Cε-Cδ - RFDR
$d_{eff}=481^{+41}_{-42}$Hz
L14Cα&Cβ-Cα - INADEQUATE
$d_{eff}=570^{+102}_{-97}$Hz
L14Cα-Cβ - RFDR
$d_{eff}=684^{+33}_{-32}$Hz
L41Cα&Cβ-Cα - INADEQUATE
$d_{eff}=557^{+76}_{-72}$Hz
L41Cα-Cβ - DARR
$d_{eff}=670^{+33}_{-33}$Hz
L41Cα-Cγ - DARR
$d_{eff}=662^{+106}_{-97}$Hz
L41Cα-Cγ - RFDR
$d_{eff}=726^{+107}_{-95}$Hz
P6Cα&Cβ-Cα - INADEQUATE
$d_{eff}=582^{+56}_{-54}$Hz
P6Cα-Cβ - DARR
$d_{eff}=612^{+51}_{-49}$Hz
P6Cα-Cβ - RFDR
$d_{eff}=649^{+38}_{-37}$Hz
P6Cα-Cγ - RFDR
$d_{eff}=702^{+99}_{-89}$Hz
P6Cα-Cδ - RFDR
$d_{eff}=655^{+147}_{-133}$Hz
P6Cγ&Cδ-Cδ - INADEQUATE
$d_{eff}=544^{+35}_{-35}$Hz
P6Cδ-Cα - RFDR
$d_{eff}=788^{+50}_{-49}$Hz
P6Cδ-Cγ - DARR
$d_{eff}=595^{+28}_{-27}$Hz
P6Cδ-Cγ - RFDR
$d_{eff}=621^{+22}_{-22}$Hz
Q15Cα&Cβ-Cα - INADEQUATE
$d_{eff}=613^{+53}_{-50}$Hz
Q15Cα-C - DARR
$d_{eff}=665^{+78}_{-74}$Hz
Q15Cα-Cβ - DARR
$d_{eff}=673^{+49}_{-48}$Hz
Q15Cγ&Cδ-Cδ - INADEQUATE
$d_{eff}=469^{+43}_{-43}$Hz
S50Cα&Cβ-Cα - INADEQUATE
$d_{eff}=606^{+68}_{-64}$Hz
S50Cα-C - RFDR
$d_{eff}=708^{+36}_{-35}$Hz
S50Cα-Cβ - DARR
$d_{eff}=675^{+70}_{-67}$Hz
$1-S/S_0$
dephasing time [ms]
$\chi^2_{red}$
Dipolar constant in Hz

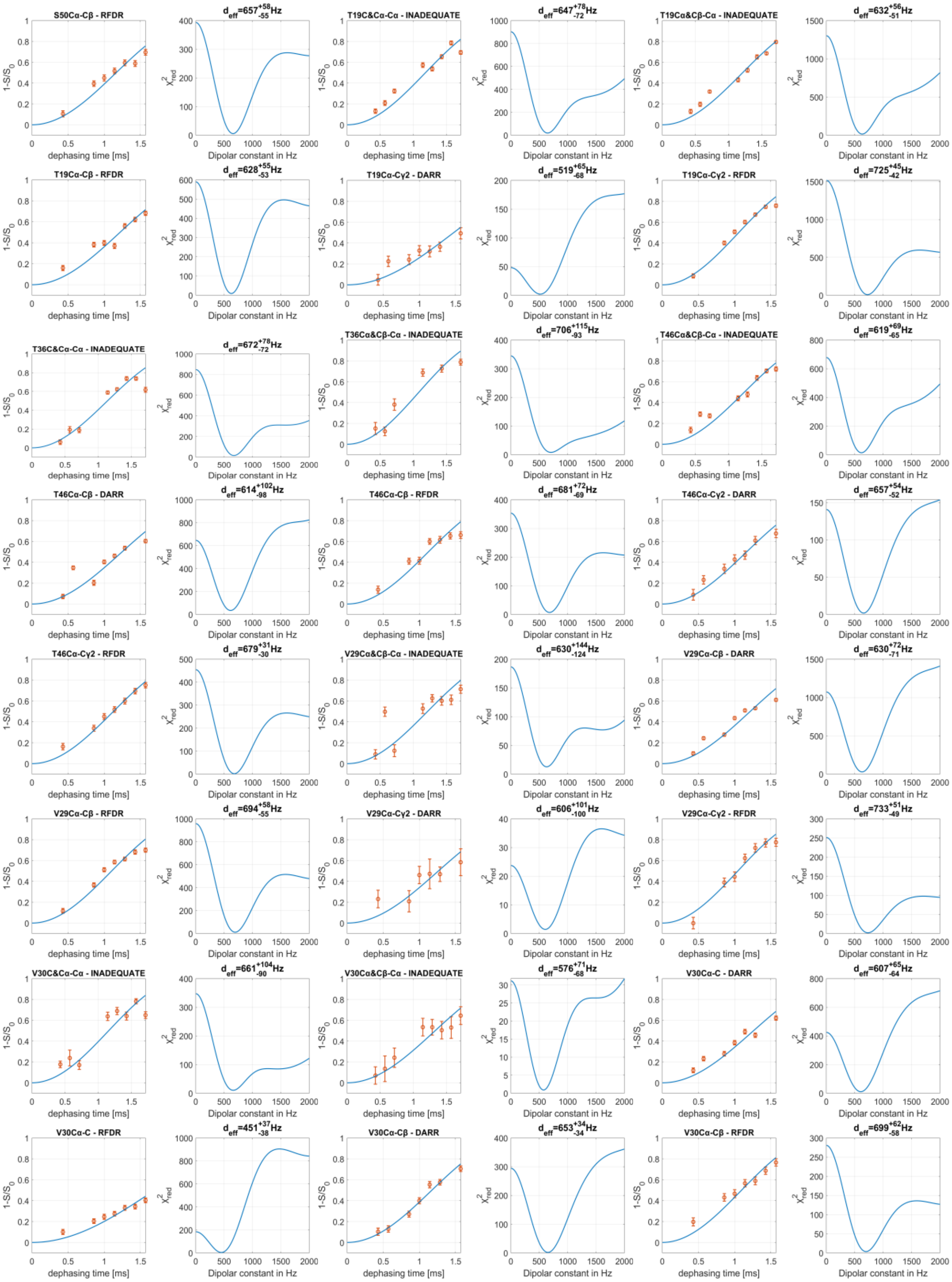
S50Cα-Cβ - RFDR
d_eff=657 +58 -55 Hz
T19C&Cα-Cα - INADEQUATE
d_eff=647 +78 -72 Hz
T19Cα&Cβ-Cα - INADEQUATE
d_eff=632 +56 -51 Hz
T19Cα-Cβ - RFDR
d_eff=628 +55 -53 Hz
T19Cα-Cγ2 - DARR
d_eff=519 +65 -68 Hz
T19Cα-Cγ2 - RFDR
d_eff=725 +45 -42 Hz
T36C&Cα-Cα - INADEQUATE
d_eff=672 +78 -72 Hz
T36Cα&Cβ-Cα - INADEQUATE
d_eff=706 +115 -93 Hz
T46Cα&Cβ-Cα - INADEQUATE
d_eff=619 +69 -65 Hz
T46Cα-Cβ - DARR
d_eff=614 +102 -98 Hz
T46Cα-Cβ - RFDR
d_eff=681 +72 -69 Hz
T46Cα-Cγ2 - DARR
d_eff=657 +54 -52 Hz
T46Cα-Cγ2 - RFDR
d_eff=679 +31 -30 Hz
V29Cα&Cβ-Cα - INADEQUATE
d_eff=630 +144 -124 Hz
V29Cα-Cβ - DARR
d_eff=630 +72 -71 Hz
V29Cα-Cβ - RFDR
d_eff=694 +58 -55 Hz
V29Cα-Cγ2 - DARR
d_eff=606 +101 -100 Hz
V29Cα-Cγ2 - RFDR
d_eff=733 +51 -49 Hz
V30C&Cα-Cα - INADEQUATE
d_eff=661 +104 -90 Hz
V30Cα&Cβ-Cα - INADEQUATE
d_eff=576 +71 -68 Hz
V30Cα-C - DARR
d_eff=607 +65 -64 Hz
V30Cα-C - RFDR
d_eff=451 +37 -38 Hz
V30Cα-Cβ - DARR
d_eff=653 +34 -34 Hz
V30Cα-Cβ - RFDR
d_eff=699 +62 -58 Hz
1-S/S0
dephasing time [ms]
χ2red
Dipolar constant in Hz

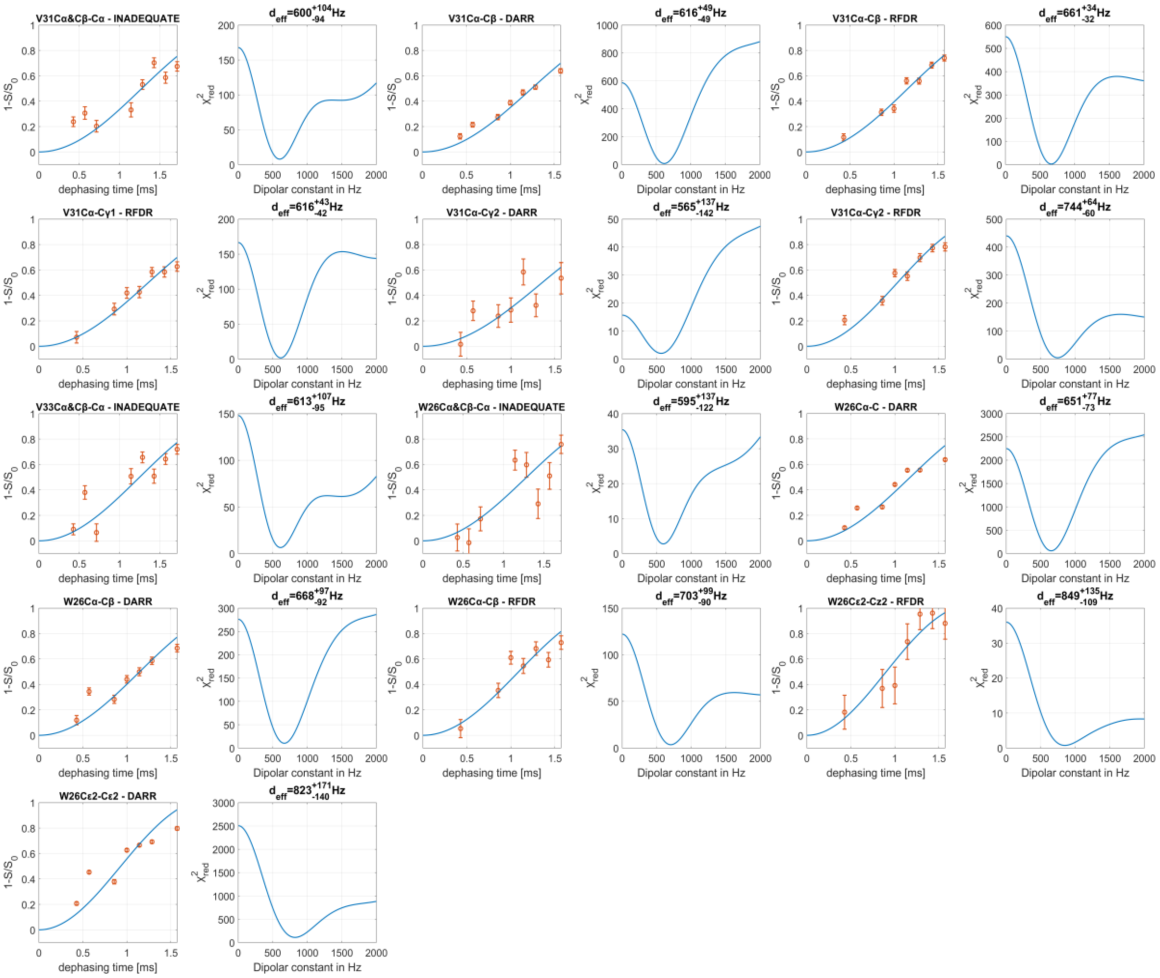

### S7 – Filtered INADEQUATE

The application of the REDOR block causes the dephasing of signals originating from carbons in close proximity to nitrogen. At the longest dephasing time that was used in our experiments, 1.714ms, approximately 80% of their signal decayed, effectively eliminating it from the spectrum.

Figure S7 shows a comparison between the same 'S' INADEQUATE-REDOR crosspeaks of threonine residues (19, 36, 46) in the fd-Y21M phage coat protein through different dephasing times. The figure shows how the signals of double-quantum Cα+Cβ crosspeaks with Cα in the acquisition dimension decay with increased dephasing times, while the crosspeaks with Cβ in the acquisition SQ dimension are not affected by the pulses, only by relaxation. The resulting edited spectrum has decreased spectral congestion while still containing information about the Cα chemical shifts encoded in the DQ dimension.

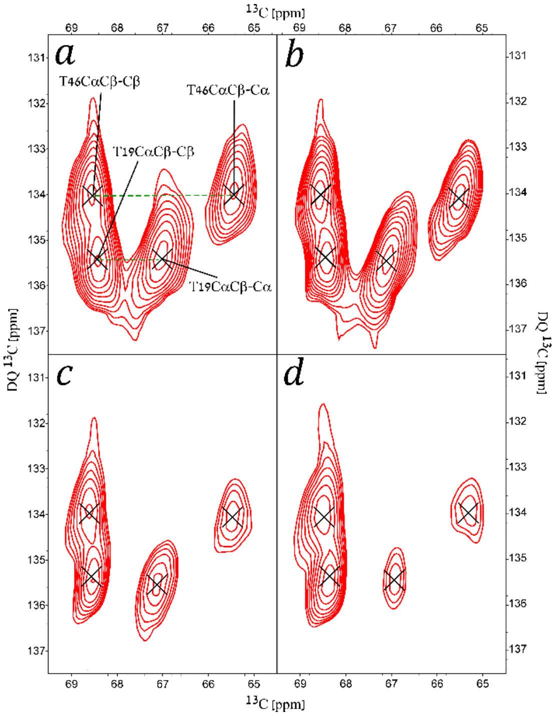

**Figure S7**. Extractions of INADEQUATE-REDOR 'S' spectra showing threonine DQ correlations with (a)-(d) corresponding to dephasing times of 429μs, 571μs, 1571μs and 1714μs, respectively. Contour levels are drawn at multiples of 1.3 with the lowest matching an SNR of 8.

In order to assess the effects of relaxation due to the addition of the REDOR block, we compared the SNR of 15 randomly selected non-ambiguous crosspeaks in both S and $S_0$ spectra. The SNR values are detailed in the following table.

| | SNR | | |
|---|---|---|---|
| Crosspeak | $S_0$ (0.429ms) | S (1.714ms) | Signal reduction |
| E2Cα&Cβ- Cβ | 25 | 16 | 36% |
| P6Cγ&Cδ- Cγ | 17 | 15 | 12% |
| K8Cγ&Cδ- Cδ | 46 | 53 | -15% |
| K8Cγ&Cδ- Cγ | 33 | 29 | 12% |
| D12Cα&Cβ- Cβ | 21 | 22 | -5% |
| I37Cγ1&Cδ1- Cδ1 | 56 | 31 | 45% |
| Q15Cα&Cβ- Cβ | 28 | 25 | 11% |
| Q15Cγ&Cδ- Cγ | 33 | 32 | 3% |
| I22Cγ1&Cδ1- Cδ1 | 41 | 37 | 10% |
| I37Cβ&Cγ2- Cβ | 31 | 15 | 52% |
| I37Cγ1&Cδ1- Cγ1 | 24 | 23 | 4% |
| I37Cβ&Cγ2- Cγ2 | 67 | 43 | 36% |
| I37Cγ1&Cδ1- Cδ1 | 56 | 31 | 45% |
| I39Cγ1&Cδ1- Cγ1 | 14 | 17 | -21% |
| I39Cγ1&Cδ1- Cδ1 | 75 | 71 | 5% |
| T46Cα&Cβ- Cβ | 52 | 59 | -13% |

On average, the change in the SNR is a decrease of 13%.
